# The oncogenic role of *Streptococcus gallolyticus subsp*. *gallolyticus* is linked to activation of multiple cancer-related signaling pathways

**DOI:** 10.1101/2023.02.17.528913

**Authors:** Ewa Pasquereau-Kotula, Giulia Nigro, Florent Dingli, Damarys Loew, Patrick Poullet, Yi Xu, Scott Kopetz, Jennifer Davis, Lucie Peduto, Catherine Robbe-Masselot, Philippe Sansonetti, Patrick Trieu-Cuot, Shaynoor Dramsi

**Affiliations:** Institut Pasteur, Université Paris Cité, CNRS UMR6047, Biology of Gram-positive Pathogens Unit, F-75015 Paris, France; Institut Pasteur, Université Paris Cité, INSERM U1224, Stroma, Inflammation and Tissue Repair Unit, F-75015 Paris, France; Institut Pasteur, Université Paris Cité, INSERM U1224, Microenvironment and Immunity Unit, F-75015 Paris, France; Institut Curie, PSL Research University, CurieCoreTech Spectrométrie de Masse Protéomique, F-75005, Paris, France; Institut Curie, Bioinformatics core facility (CUBIC), INSERM U900, PSL Research University, Mines Paris Tech, F-75005, Paris, France; Center for Infectious and Inflammatory Diseases, Institute of Biosciences and Technology, Texas A&M Health Science Center; Department of Microbial Pathogenesis and Immunology, School of Medicine, Texas A&M Health Science Center; Department of Microbiology and Molecular Genetics, University of Texas Health Science Center, Houston, Texas, United States of America; Department of Gastrointestinal Medical Oncology, the University of Texas MD Anderson Cancer Center, Houston, Texas, United States of America; Univ. Lille, CNRS, UMR8576 - UGSF - Unité de Glycobiologie Structurale et Fonctionnelle, F-59000, Lille, France; Institut Pasteur, Unité de Pathogénie Microbienne Moléculaire, INSERM U1202, and College de France, F-75005, Paris, France

**Author notes:** University of Kansas, Kansas City, Kansas (Jennifer Davis).

**Keywords:** *Streptococcus gallolyticus*, colorectal cancer, proteome, phosphoproteome

## Abstract

*Streptococcus gallolyticus subsp. gallolyticus (SGG)*, an opportunistic gram-positive pathogen responsible for septicemia and endocarditis in the elderly, is often associated with colon cancer (CRC). In this work, we investigated the oncogenic role of *SGG* strain UCN34 using the azoxymethane (AOM)-induced CRC model *in vivo*, organoid formation *ex vivo* and proteomic and phosphoproteomic analyses from murine colons. We showed that *SGG* UCN34 accelerates colon tumor development in the murine CRC model. Full proteome and phosphoproteome analysis of murine colons chronically colonized by *SGG* UCN34 or the closely related non-pathogenic *S. gallolyticus subsp. macedonicus* (*SGM*) revealed that 164 proteins and 725 phosphorylation sites were differentially regulated following colonization by *SGG* UCN34. Ingenuity Pathway Analysis (IPA) indicates a pro-tumoral shift specifically induced following colonization by *SGG* UCN34, as most proteins and phosphoproteins identified were associated with digestive cancer. Comprehensive analysis of the altered phosphoproteins using ROMA software revealed significantly elevated activities in several cancer hallmark pathways affecting tumoral cells and their microenvironment, i.e. MAPK (ERK, JNK and p38), mTOR and integrin/ILK/actin signaling, in *SGG* UCN34 colonized colon. Importantly, analysis of protein arrays of human colon tumors colonized with *SGG* showed up-regulation of PI3K/Akt/mTOR and MAPK pathways, providing clinical relevance to our findings. To test *SGG*’s capacity to induce pre-cancerous transformation of the murine colonic epithelium, we grew *ex vivo* organoids which revealed unusual structures with compact morphology following exposure to *SGG*. Taken together, our results reveal that the oncogenic role of *SGG* UCN34 is associated with activation of multiple cancer-related signaling pathways.

**Author Summary:** Colorectal cancer is the third most common cause of cancer mortality worldwide. The colon is a very singular organ, colonized by a vast and complex community of microorganisms, known as the gut microbiota. Strong evidence supports a role of the microbiota in colon cancer development. *Streptococcus gallolyticus* subsp. *gallolyticus* (*SGG*), a gut commensal, was one of the first bacteria to be associated with colorectal cancer. A better understanding of the role of *SGG* in colon cancer development is critical to developing novel strategies to improve clinical diagnosis and treatment of this disease. Here, using a global proteomic analysis of mouse colonic tissue colonized by *SGG*, we show that over 90% of the proteins with altered levels are involved in cancer. *SGG* colonization promotes autocrine and paracrine pro-tumor signals contributing to transformation of the colonic epithelium but also modifying the stromal microenvironment, which in turn sustains tumor development. Importantly, two tumor hallmark pathways (PI3K/Akt/mTOR and MAPK) identified in our mouse model were also found in human colon tumor biopsies colonized by *SGG*, strengthening the clinical relevance to our study.

## Introduction

Colorectal cancer (CRC) is the third most diagnosed cancer worldwide with 1.9 million new cases detected in 2020 [1]. CRC develops in patients over many years through the accumulation of genetic and epigenetic alterations. Many intrinsic or extrinsic factors can influence the initiation and/or development of CRC, complicating the study of its etiology.

The colon is composed of several cellular layers, including an epithelial layer and a subepithelial lamina propria containing abundant stromal cells, which are required for intestinal homeostasis [2]. Both epithelial cells and their stromal microenvironment play a role in CRC development [3]. The colon is a very singular organ colonized by a vast and complex community of microorganisms known as the gut microbiota. It is composed of over 500 species totaling approximately 10^13^ bacteria. Accumulating evidence supports the role of the microbiota in CRC development [4–8]. A few bacterial species have been identified as playing a role in colorectal carcinogenesis including: *Streptococcus gallolyticus* subsp. *gallolyticus* (*SGG*) *Fusobacterium nucleatum*, enterotoxigenic *Bacteroides fragilis*, colibactin-producing *Escherichia coli, Parvimonas micra* and *Clostridium septicum* [9]. Understanding the role of *SGG* in CRC is critical to developing novel strategies to improve clinical diagnosis and treatment of this disease.

*SGG*, formerly known as *Streptococcus bovis* biotype I, is one of the first bacteria to be associated with CRC [10,11]. This association, ranging from 47 to 85%, has been confirmed by several epidemiological studies [12–18]. Most of these studies were performed on a selected cohort of patients with a history of *SGG* invasive infection (bacteremia and/or endocarditis). We studied *SGG* prevalence in control or CRC-patients, showing that *SGG* was detected more frequently in the stools of patients with adenocarcinomas (50%) as compared to early-adenoma patients (30%) [19].

*S. gallolyticus* has been subdivided into three subspecies, subsp. *gallolyticus* (*SGG*), subsp. *pasteurianus* (*SGP*), and subsp. *macedonicus (SGM)* (**Fig. S1A**). Among them, only *SGG* is associated with CRC, suggesting that *SGG*-specific attributes contribute to this association. *SGM*, the genetically closest non-pathogenic species, was used in this study as a control bacterium. Whether *SGG* is a driver and/or a passenger bacterium in CRC has been an open question for decades (reviewed in [20]). In favor of the passenger model, our group and others previously showed that tumor-associated conditions provide *SGG* UCN34 with the ideal environment to proliferate [21,22]. We showed that this colonization advantage was linked to the production of a bacteriocin enabling *SGG* to kill closely related gut microbiota bacteria [22]. In favor of the driver model, Kumar et al. showed that *SGG* strain TX20005 accelerates tumor growth through activation of the Wnt/β-catenin signaling pathway, leading to increased cell proliferation [23,24]. The same group reported more recently that a type VII secretion system of *SGG* TX20005 contributes to the development of colonic tumors [25]. Lastly, it was also suggested that *SGG* ATCCBAA-2069 drives tumor formation by modulating anti-tumor-immunity in a colitis-associated CRC model [26].

In this work, we tested the oncogenic potential of *SGG* UCN34 using 2 different murine CRC models: (i) the AOM-induced CRC model in A/J mice and (ii) the genetic APC^Min/+^ model. Azoxymethane (AOM) is an active derivative of dimethyl hydrazine, an azide compound inducing DNA mutations in colon cells [27]. The AOM-induced mouse model reflects the human sporadic CRC events with adenoma to carcinoma progression which are largely influenced by gut microbiota composition. As a second CRC model, we used a genetic APC^Min/+^ model. Multiple intestinal neoplasia (Min) mice, containing a point mutation in the *Adenomatous polyposis coli* (APC) gene in a C57BL/6 background (C57BL/6 APC^Min/+^ = APC^Min/+^) [28], develop numerous adenomas in the small intestine and was the first model used to demonstrate the involvement of the APC gene in intestinal tumorigenesis [29,30].

We show that *SGG* UCN34 can accelerate tumor development in the AOM-induced CRC model compared to mice colonized with control *SGM*. Analysis of the global proteome and phosphoproteome from murine colon tissue colonized by *SGG* UCN34 or control *SGM* over 12 weeks revealed that *SGG* UCN34 promote autocrine and paracrine pro-tumor signals. Most of the proteins and phosphoproteins regulated by *SGG* UCN34 were associated with digestive cancer. In depth bioinformatic analyses using ROMA software revealed that the pro-oncogenic role of *SGG* UCN34 is associated with activation of several cancer hallmark pathways in tumor cells and their stromal microenvironment, i.e. MAPK (ERK, JNK and p38), mTOR and integrin/ILK/actin signaling. In line with these results, we show here for the first time that *SGG* UCN34 contributes to pre-cancerous transformation of the colon epithelium using an *ex-vivo* organoid model. Finally, protein analysis of human colon tumors colonized with *SGG* revealed up-regulation of PI3K/Akt/mTOR and MAPK pathways, providing clinical relevance to our findings in a murine model of CRC.

## Results

### 1. *SGG* UCN34 induces acceleration of tumorigenesis in AOM-induced CRC model

We tested the oncogenic potential of *SGG* UCN34 in the AOM-induced CRC model in A/J mice as summarized in **Fig. 1A**. We compared groups of mice infected for 12 weeks with *SGG* UCN34 or the non-pathogenic control bacterium *SGM*. A third group of mice, annotated NT, received PBS by oral gavage. We showed that both *SGG* UCN34 and *SGM* were able to colonize the murine colon at comparable levels through the duration of the experiment (**Fig. 1B**). *SGM* colonization remained very stable over the 12 weeks while *SGG* UCN34 showed only a slight decrease after week 8. After 12 weeks of colonization, mice were sacrificed, and colons dissected for tumor counts. We found that mice colonized by *SGG* UCN34 displayed between 1 and 8 tumors (average of 4 tumors/mouse), mostly in the distal and middle part of the colon, whereas in the control *SGM* group only 2 mice exhibited 1 tumor. In the NT group, only 1 mouse showed 1 tumor (**Fig. 1C**). In addition, the few tumors recorded in the *SGM* and NT control groups were very small in size compared to those observed with *SGG* UCN34 (**Fig. 1D**). Histopathological analyses of the colonic tumors indicate that most of the tumors induced by *SGG* UCN34 were low-grade adenomas (**Fig. 1E, Fig. S1A**), while low-grade dysplasia and low-grade adenomas were observed in NT and *SGM* groups (**Fig. 1E, Fig. S1**A).

**Figure 1.**
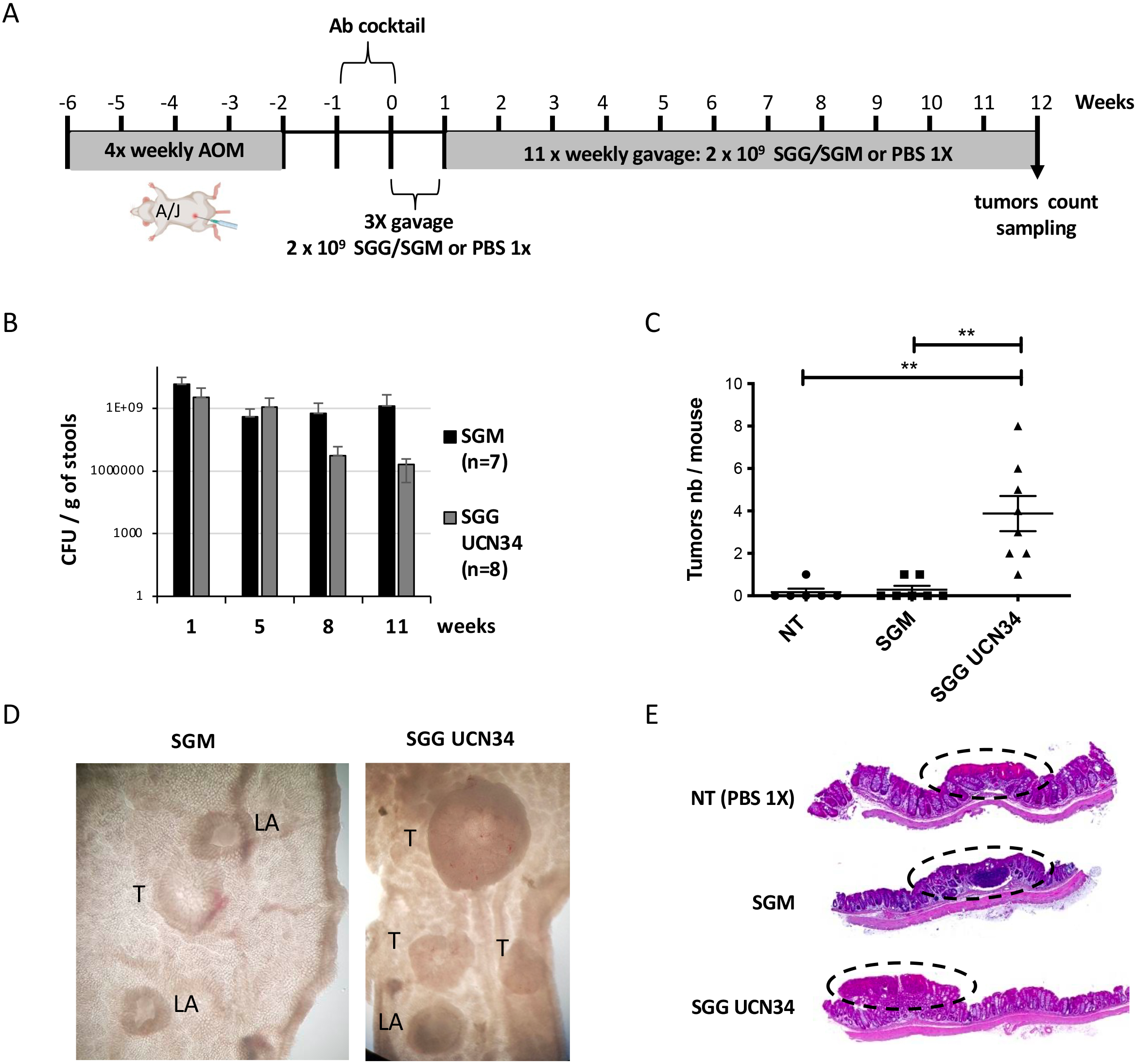
*SGG* UCN34 induces acceleration of tumorigenesis in AOM-induced CRC model. The effect of *SGG* UCN34 vs *SGM* on the development of colonic tumors was examined in an AOM-induced CRC model, as shown in **A**. **B.** CFU (colony forming units) per g of stools. Fecal materials were collected at different time points (1, 5, 8 and 11 weeks), homogenized, and serial dilutions plated onto Enterococcus Selective Agar plates to count *SGG* UCN34 and *SGM* bacteria. **C.** Sum of tumors per mouse. Macroscopic tumors were evaluated by an experimented observer. ns, not significant; *, *p* < 0.05; **, *p* < 0.01; Mann-Whitney test. **D**. Representative pictures of mouse colon after dissection for *SGM* and *SGG* UCN34 groups (T = tumor, LA = lymphoid aggregate). **E.** Representative histological (H&E) sections of colon tumors found in AOM-induced CRC mouse model for each experimental group: NT (low grade dysplasia), *SGM* (low grade dysplasia) and *SGG* UCN34 (low grade adenoma).

We compared the oncogenic effect of *SGG* UCN34 with other *SGG* isolates distant from UCN34. We chose the USA isolate TX20005 previously shown to accelerate tumor formation in the AOM-induced CRC model [23–25]. Mice colonized over 12 weeks with *SGG* UCN34 or TX20005 exhibited a higher number of tumors as compared to control groups NT and *SGM* (**Fig. S1B**). There was no significant difference in term of tumor count between the two UCN34 and TX20005 *SGG* isolates (**Fig. S1B**).

We also tested the oncogenic potential of *SGG* UCN34 in the APC^Min/+^ mouse model (**Fig. S2A)**. As shown in **Fig. S2B**, mice colonized with *SGG* UCN34 exhibited a slightly increased numbers of adenomas from small intestine that were also larger compared to mice colonized with control *SGM* bacteria, but these differences were not statistically significant. We hypothesize that this may be due to a lower capacity of colonization of the small intestine by *SGG* UCN34 (400X times less) as compared to the colon (**Fig. S2C**).

### 2. *SGG* UCN34 induces multiple pro-tumoral changes in murine colon epithelium

In order to decipher the molecular mechanisms underlying *SGG*-induced tumorigenic transformation, full proteome and phosphoproteome analyses were carried out on whole protein extracts from macroscopically tumor-free colon tissue of mice colonized by *SGG* UCN34 or *SGM* as well as from colonic tumors of mice exposed to *SGG* UCN34 (**Fig. 2A**). Phosphoproteome analysis was performed by using proteome samples and phosphopeptide enrichment followed by label-free quantitative mass spectrometry.

**Figure 2.**
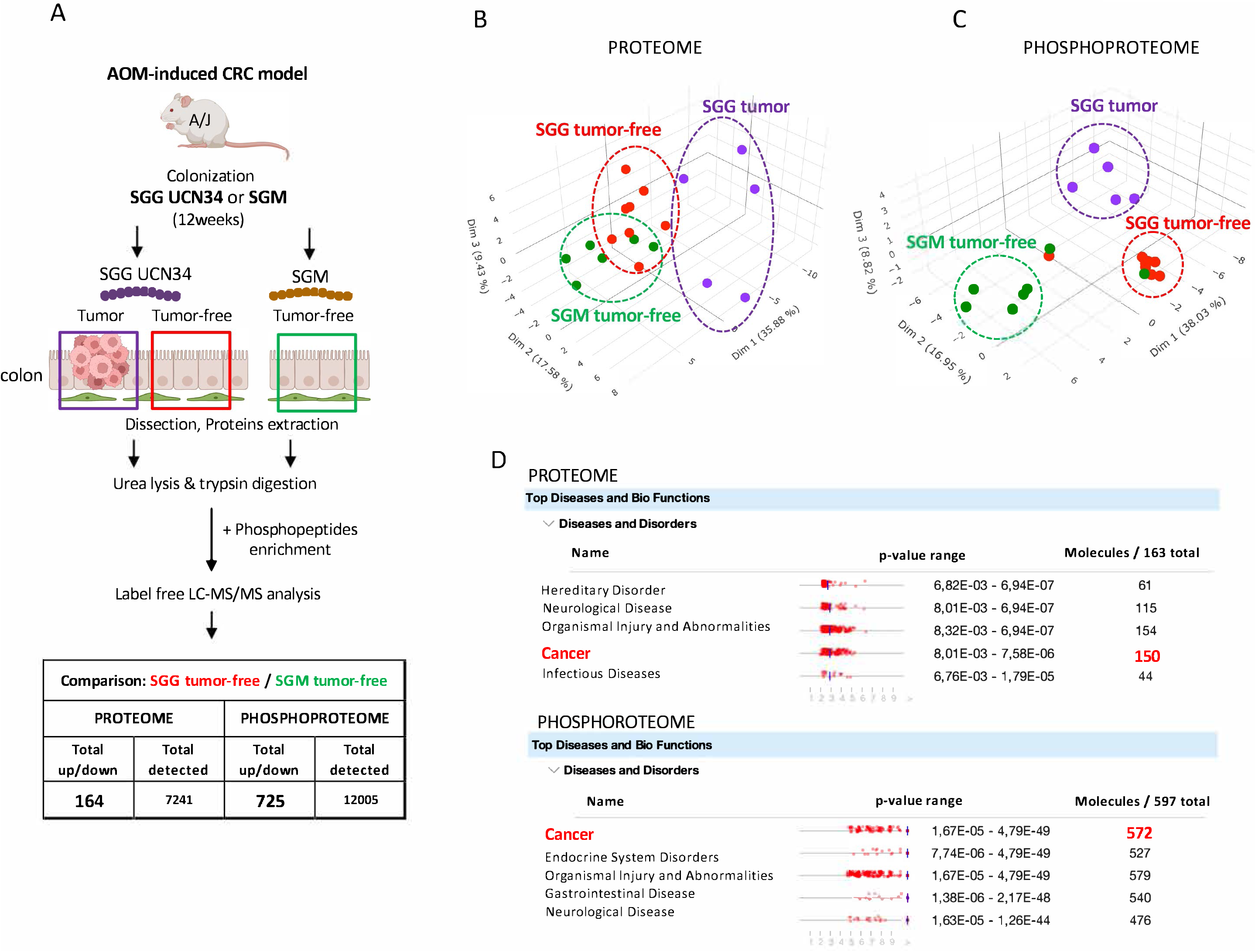
Proteome and phosphoproteome analyses on colonic tissue colonized by *SGG* UCN34 or *SGM* over 12 weeks reveals a pro-tumoral shift specific to *SGG* UCN34. **A.** Sample preparation for proteome and phosphoproteome LC-MS analysis. All samples were collected at the end of the AOM-induced CRC model. Tissue samples were from macroscopically tumor-free colon sections for *SGM* and *SGG* UCN34 groups and from colon tumors found in this experiment only for the *SGG* UCN34 group. Proteins were extracted by urea lysis and trypsin digestion. For phosphoproteome MS analysis a phosphopeptide enrichment step was added. All samples were processed by label free LC-MS/MS analysis. **B.** Principal Component Analysis (PCA) of proteome samples in 3D representation. **C.** PCA of phosphoproteome samples in 3D representation. Proteome and phosphoproteome analyses were done using myProMS web server [56]. **D**. Top Disease and Bio Functions (IPA) of proteome and phosphoproteome changes between *SGG* UCN34 and *SGM*. The Diseases & Functions Analysis identified biological functions and/or diseases that were most significant from the data set. Data sets used in this analysis were proteins or phosphoproteins differentially expressed between *SGG* UCN34 and *SGM* detected in macroscopically tumor-free colonic tissue. Molecules indicate the number of proteins or phosphoproteins associated with indicated diseases and disorders. A right-tailed Fisher’s Exact Test was used to calculate a *p*-value determining the probability that each biological function and/or disease assigned to that data set is due to chance alone.

The proteome of tumor-free colon segments in mice colonized with *SGG* UCN34 or *SGM* revealed that *SGG* UCN34 induces significant changes in the levels of 164 proteins as compared to the control *SGM* group (**Fig. 2A**, **Sup Table 1**). Most of these protein levels were down-regulated (129) and 35 were up-regulated. Even stronger changes were observed in the phosphoproteome with as many as 725 phosphosites (642 proteins) at significantly different levels between *SGG* UCN34 vs *SGM:* 325 sites on 299 proteins up- and 400 sites on 343 proteins down-regulated (**Fig. 2A, Sup Table 2**).

Principal Component Analyses (PCA) of all samples based on protein (**Fig. 2B**) and phosphosite abundance (**Fig. 2C**) revealed 3 separate clusters matching the phenotypic observations (*SGM* tumor-free, *SGG* tumor-free, *SGG* tumor). As expected, the protein content of *SGG*-induced tumors differs from that of the adjacent macroscopically healthy tissue. Importantly, *SGG* UCN34 induced specific proteomic and phosphoproteomic changes in macroscopically tumor-free tissue compared to *SGM*-treated tumor-free tissue indicating that *SGG* UCN34 modulates the proteome and phosphoproteome landscapes of the murine colon epithelium before any macroscopically visible changes. These results strongly suggest that *SGG* UCN34 is not a silent colon inhabitant.

To investigate the molecular functions altered by *SGG* UCN34, Ingenuity Pathway Analysis (IPA, Qiagen) was applied to our datasets. We found that the differentially expressed proteins and phosphosites in tumor-free *SGG* UCN34 vs *SGM* were involved mainly in cancer disease (**Fig. 2D**). As many as i) 150 proteins out of 163 and ii) 572 phosphoproteins out of 597 were predicted to be associated with cancer development.

This is the first demonstration that *SGG* UCN34 can induce a clear and massive pro-tumoral shift at the proteomic and phosphoproteomic levels in mice colons. Our results further indicate that *SGG* UCN34 has an impact not only on existing tumoral lesions, but also on adjacent tumor-free segments of the colon.

### 3. *SGG* UCN34 activates MAPK cascades, mTOR and integrin/ILK/actin cytoskeleton pathways *in vivo*

To decipher the signaling pathways altered by *SGG* UCN34, we processed our phosphoproteomic data sets with the in-house built ROMA (Representation and quantification Of Module Activities) software, a gene-set-based quantification algorithm [31] integrating several databases and the commercially available IPA (Ingenuity Pathway Analysis) software. Comparison of the three groups, namely *SGG* UCN34 tumor-free, *SGM* tumor-free and *SGG* UCN34 tumor, revealed 83 pathways significantly regulated (evidenced by two-tailed *t*-tests of activity scores; p-value ≤0.05) including 654 different genes (**Fig. 3A, Sup Table S3**). Among these 83 pathways, 34 were up-regulated by *SGG* UCN34 and 49 pathways were down-regulated. The top 10 up-regulated pathways were the following: epidermal growth factor (EGF), integrin, platelet derived growth factor (PDGF), PYK2, melanoma, p38, mitogen activated protein kinases (MAPK) family signaling cascades, target of rapamycin (mTOR) signaling, regulation of Ras by GTPase activating proteins (GAPs), signaling by fibroblast growth factor receptor 2 (FGFR2) (**Sup Table S3**). Five of these, EGF, PDGF, p38, GAPs and FGFR2, converge in MAPK family signaling cascades. Interestingly, although expression of EGFR and FGFR2 is more ubiquitous, PDGFRA (one of the major phosphoproteins identified in the PDGF pathway, **Sup Table. S5**) is exclusively expressed by stromal cells in normal intestine as well as in colorectal cancer [32,33] (**Fig. S6A-B**), suggesting involvement of the stromal microenvironment. Apart from the MAPK cascades, the results point out activation of mTOR signaling pathway as well as integrin signaling. The top 10 down-regulated pathways were the following: adherens junction, insulin signaling, vascular smooth muscle contraction, protein tyrosine phosphatase 1B (PTP1B), protein tyrosine phosphatase-2 (SHP2) pathway, AUF1 (hnRNP D0) which binds and destabilizes mRNA, Feline McDonough Sarcoma-like tyrosine kinase (FLT3) signaling, RUNX2 which regulates bone development, transcriptional regulation by TP53, Fc epsilon receptor 1 (FCER1) signaling (**Sup Table S3**). Thus, most down-regulated pathway were linked to actin cytoskeleton modulation, through adherens junction, cell junction organization, focal adhesion and cell-cell communication (**Sup Table S3**).

**Figure 3.**
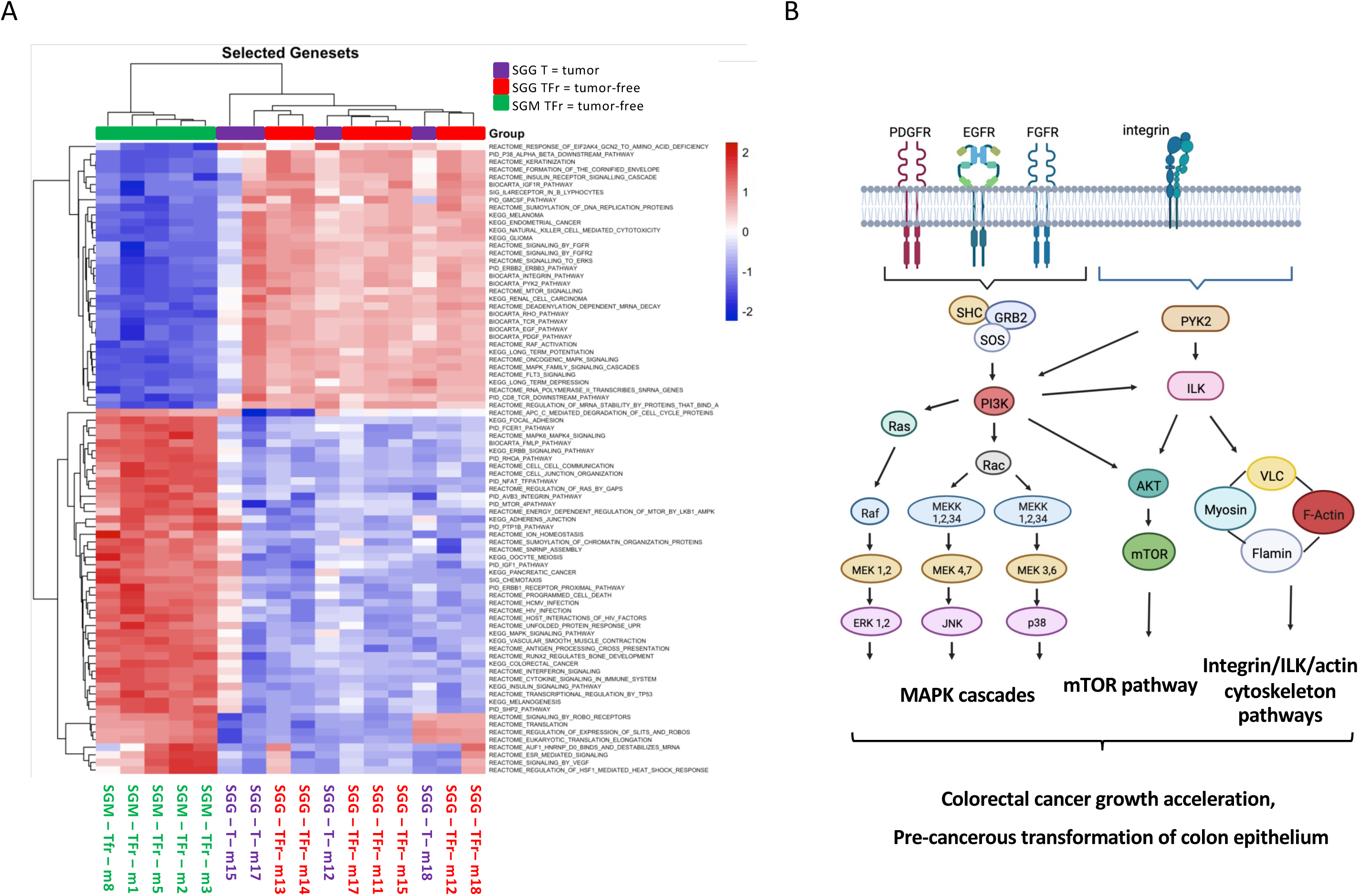
ROMA analysis reveals multiple cancer-related pathways altered by *SGG* UCN34 in comparison to *SGM*. **A.** Heat map view of up and downregulated pathways between three groups: *SGM* tumor-free, *SGG* UCN34 healthy/tumor-free and *SGG* UCN34 tumors. **B.** Schematic diagram of *SGG* UCN34-induced activation of MAPK cascades, mTOR signaling and integrin/ILK/actin cytoskeleton pathways. ROMA pathway analysis together with IPA indicates activators of MAPK cascades by UCN34. Apart from the MAPK cascades we can point out activation of the mTOR signaling pathway as well as integrin/ILK/actin cytoskeleton signaling. Created with BioRender.com. ROMA = Representation and quantification Of Module Activities.

IPA analysis of phosphoproteomic changes between *SGG* UCN34 tumor-free and *SGM* tumor-free also revealed activation of several cancer-related pathways. Amongst the top 10 selected by activation prediction score (positive z-score) were: Integrin-linked kinase (ILK) signaling, BRCA1 in DNA damage response, Dilated Cardiomyopathy Signaling Pathway, Protein Kinase A Signaling, Actin Cytoskeleton Signaling, AMPK signaling, Epithelial Adherens Junction Signaling, Oxytocin Signaling Pathway, Integrin Signaling and CNTF Signaling (**Fig. S3)**.

Both IPA and ROMA analyses pinpointed three highly interconnected signaling pathways namely i) ERK/MAPK ii) mTOR (**Fig. S4**) and iii) integrin/ILK/actin cytoskeleton (**Fig. S5**).

Taken together, our results suggest that *SGG* UCN34 can interfere with several pro-tumoral signaling pathways, such as MAPK cascades, mTOR and integrin/ILK/actin cytoskeleton pathways (**Fig. 3B)** which in turn can contribute to colorectal cancer growth acceleration as well as pre-cancerous transformation of the colon epithelium.

### 4. Protein Analysis of human colon tumors enriched for *SGG* reveals up-regulation of PI3K/Akt/mTOR and MAPK pathways

To investigate whether the signalling pathways activated by *SGG* in the AJ/AOM murine model of CRC were relevant in human disease, we compared the protein content of human colon tumor biopsies colonized with *SGG* (n=32) vs control colon tumor biopsies negative for *SGG* (n=29) [23] using RPPA (Reverse Phase Protein Array) [34]. In total, 52 proteins and/or phosphosites were differentially regulated between *SGG*-enriched human colon tumors vs *SGG*-negative ones. Among them, 16 proteins and 11 phosphosites were up-regulated and 17 proteins and 8 phosphosites were down-regulated (**Fig. 4**, **Sup Table S4**). Interestingly the majority of the up-regulated proteins/phosphosites belong to two pathways: (i) PI3K/AKT/mTOR (Akt, Akt pS473, GSK3 p89, GSK3αβ, GSK3αβ PS21, PI3K p85, PTEN, Src pY527, TSC1, PDK1, Lck, Smad2, β-Catenin) and (ii) MAPK (p38, p38 pT180pY182, JNK2, Syk, PKCa pS657, PKCβII pS660).

**Figure 4.**
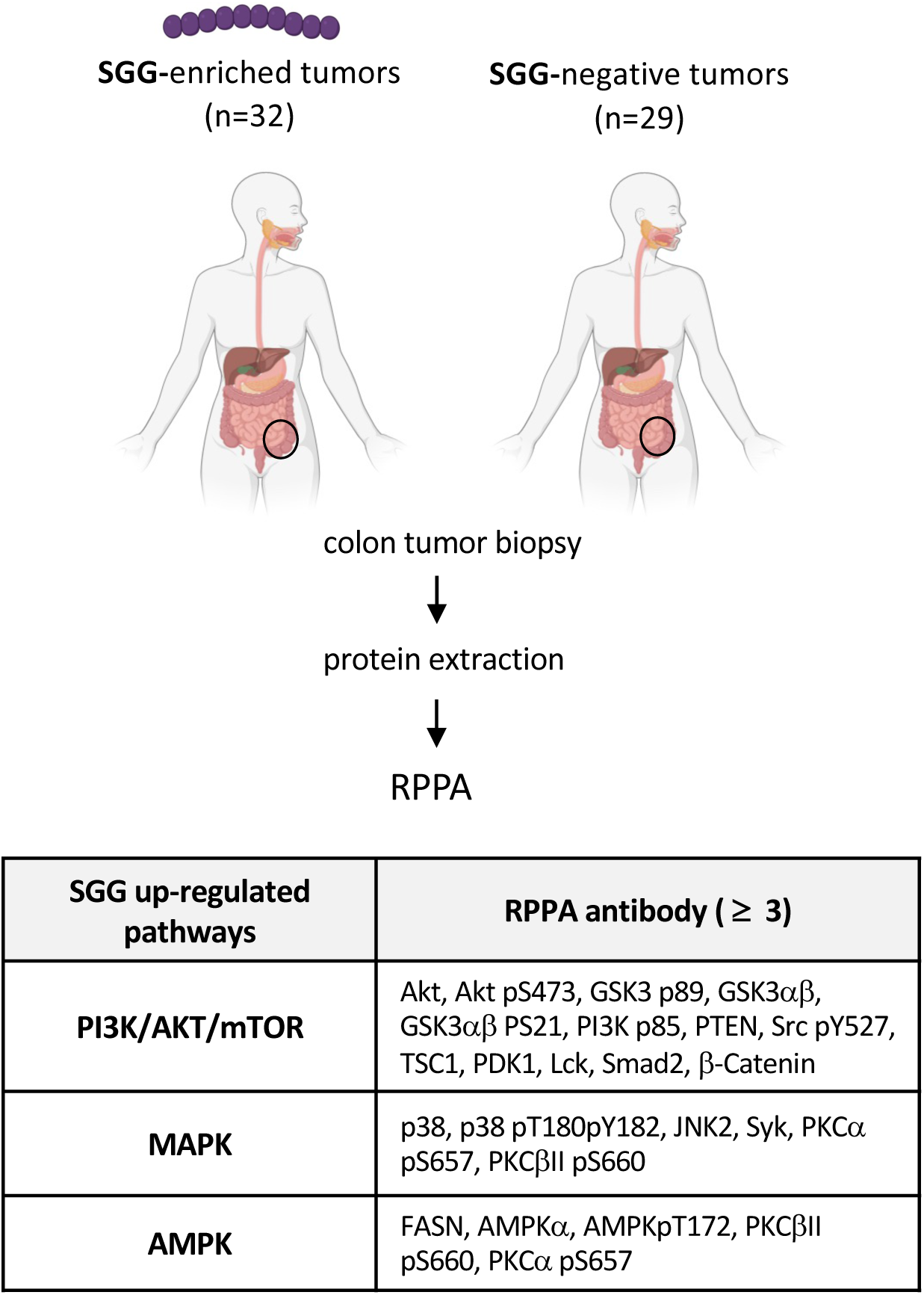
*SGG* induces up-regulation of PI3K/Akt/mTOR, MAPK and AMPK pathways in human colon tumors. Proteins were extracted from human colon biopsies enriched for *SGG* (n=32) vs negative ones (n=29) and subjected to RPPA. The table present up-regulated by *SGG* pathways for which at least 3 RPPA targeted antibody were attributed. RPPA = Reverse Phase Protein Array.

For the down-regulated proteins and/or phosphosites, no clear grouping into specific pathways could be found, and this included androgen receptor signaling, DNA damage repair, cell cycle, ERa signaling, PI3K/AKT/mTOR, MAPK (**Sup Table S4**).

### 5. *SGG* UCN34 induces formation of abnormal *ex vivo* organoids with compact morphology

Lastly, we asked whether the observed pro-tumoral proteomic and phosphoproteomic shifts in tumor-free sections of murine colon exposed to *SGG* UCN34 has biological consequences on this tissue. In other words, if *SGG* UCN34 could initiate pre-cancerous transformation of murine colon epithelium. One of the major upregulated pathways upon *SGG* UCN34 infection was PDGF. Within this pathway we identified phosphorylated form of PDGFRA (pS767; **Sup Table S5**), a receptor strongly involved in the stroma-cancer cells crosstalk [35]. By analysis of single cells RNAseq data from human colorectal cancer and normal adjacent tissue [32] (**Fig. S6**), we observed that *Pdgfra* is mostly expressed by colonic stromal cells positive for podoplanin (*Pdpn*^+^). These stromal cells are the main producers of bone morphogenetic protein (BMP) antagonists such as Gremlin 1 (*Grem1*) involved in stem cells maintenance and tumorigenesis [36–38] (**Fig. S6B**). By histology, the intestinal Grem1 producing stromal niche, which support intestinal stem cells, can be further identified by costaining Pdpn^+^ cells for CD34 [39]. Stromal cells positive for Pdpn and CD34 are present in normal colon as well as in colon cancer [39,32] (**Fig. S6B-C**). In histological sections of macroscopically tumor-free regions, we observed increased frequency of Pdpn^+^CD34^+^ stromal cells in the *SGG* UCN34 group compared to the *SGM* group (**Fig. 5A**), consistent with an early involvement of the colonic stromal microenvironment in SGG-induced tumorigenesis.

**Figure 5.**
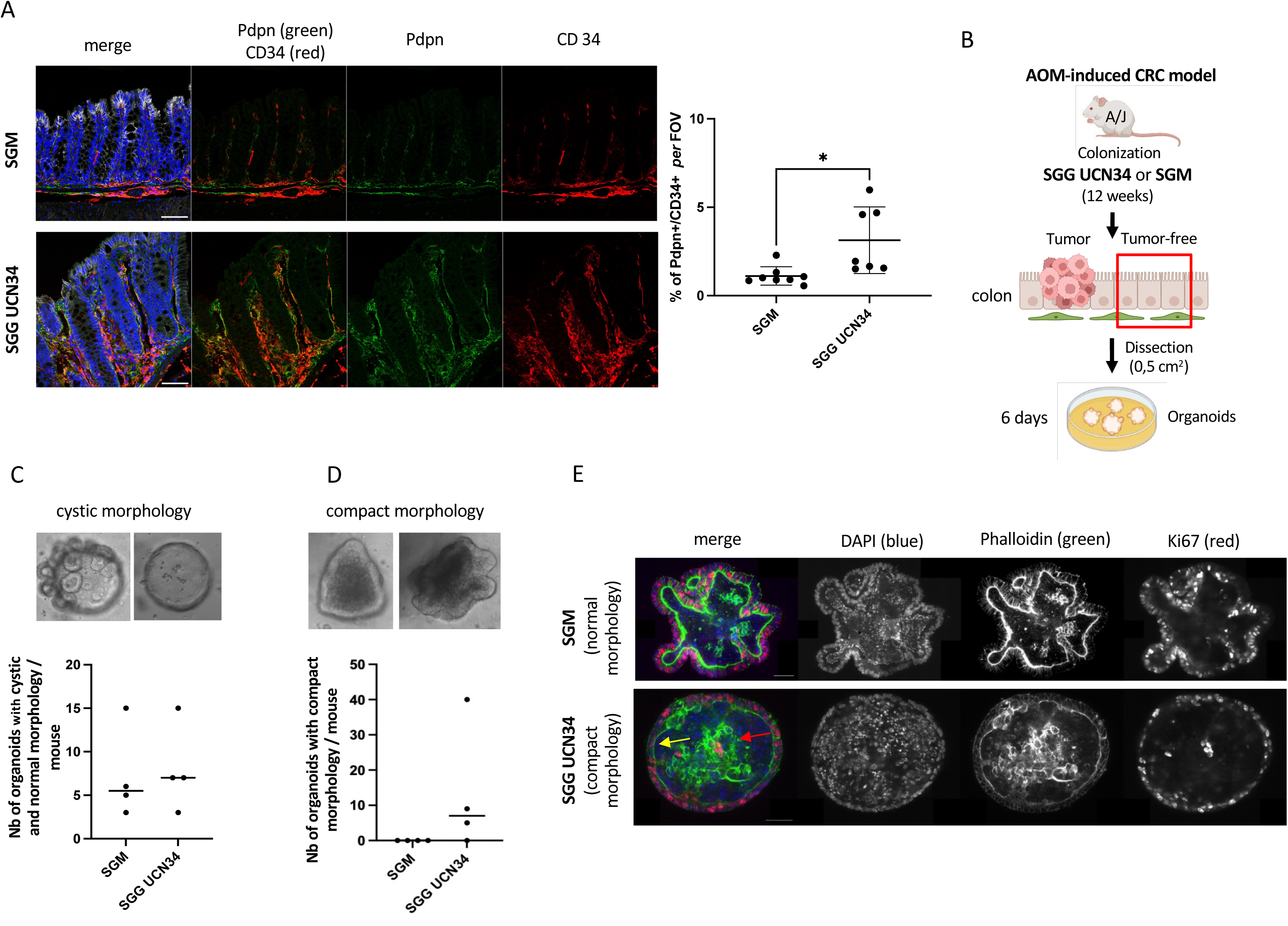
*SGG* UCN34 induces pre-cancerous transformation of murin colon tissue. **A.** Immunofluorescence analysis of Pdpn (green) and CD34 (red) in sections of colonic tumor-free regions of mice colonized with *SGM* (top) or *SGG* UCN34 (bottom) (DAPI/nuclei in blue and E-cadherin in grey). Scale bar: 50 μm. Images were acquired with SP8 microscope (Leica) and a 40X objective. Graph presents the percentage of colocalization of Pdpn and CD34 signals per field of view (FOV). The quantification was done by Imaris (version8) software on *SGM* (n=8) and *SGG* UCN34 (n=7) colonized mice, using at least 6 images per mouse and showing 1.1% vs 3.1% of Pdpn+CD34+ signal respectively (*, t-test p=0.01). **B**. Experimental design of *ex vivo* organoid formation originating from macroscopically tumor-free sections of colon tissue colonized by *SGG* UCN34 or *SGM* over 12 weeks in the AOM-induced CRC model. **C.** The number of organoids with normal or cystic morphology defined as colonospheres or colonoids with a central empty lumen and polarized epithelium (actin staining). **D.** The number of organoids with compact morphology defined as colonospheres or colonoids with a lumen full of cells and depolarized and restructured actin in the epithelium. **E.** Colored confocal pictures show DAPI/nuclei labeling (blue), Phalloidin/actin (green) and Ki67/proliferation marker (red). Each time we show one section of the middle of organoid. All images were acquired with Opterra microscope (Bruker) at 63X objective and analyzed by Fiji software. Scale bar: 50 μm.

To determine whether intestinal epithelial cells were intrinsically modified in their capacity to proliferate and differentiate, we examined *ex vivo* organoid formation [40]. Normally, organoid formation from murine colonic crypts requires supplementation of several niche factors (Wnt, R-Spondin, EGF, Noggin) to maintain their stemness and proliferation status [41]. We hypothesized that pre-cancerous tissue transformation of the colonic tissue in the group of mice colonized with *SGG* could allow organoid formation without addition of niche factors.

Development of *ex vivo* organoids originating from cells obtained from tumor-free colon tissue colonized by *SGG* UCN34 or *SGM* over 12 weeks in the AOM-induced CRC model was tested (**Fig. 5B**). After 6 days of culture in the absence of the four niche factors (Wnt, R-Spondin, EGF, Noggin), we evaluated the presence or absence of organoids and classified them according to their morphology. In both groups, we observed a similar number of organoids that we classified as normal or cystic, characterized by budding crypts or round structures respectively, composed of a monolayer of epithelial cells with a central lumen containing some dead cells [40] (**Fig. 5C**). Strikingly, we found that only *SGG* UCN34 induced the formation of numerous organoids with a different morphology, defined as compact (**Fig. 5D**). These compact organoids are characterized by their heterogeneous shape, and a filled lumen. The epithelium of compact organoids appeared depolarized and unstructured compared to the polarized and organized epithelium observed in the normal or cystic organoids, as highlighted by phalloidin staining (yellow narrow, **Fig. 5E**). Staining of these abnormal organoids induced by *SGG* with the proliferative Ki-67 marker revealed the presence of proliferative cells within the lumen (red arrow, **Fig. 5E**). Other examples of organoids with normal and cystic morphology and organoids with compact morphology are presented in **Fig. S7.**

Taken together these results show for the first time that *SGG* UCN34 contributes to early (macroscopically not visible) pre-cancerous transformation of murine colon epithelium.

## Discussion

A better understanding of the contributions of *Streptococcus galloyticus* subsp. *gallolyticus* (*SGG*) to colorectal cancer (CRC) acceleration is critical to developing novel strategies to improve clinical diagnosis and treatment of this disease.

In this work, we have shown that *SGG* strain UCN34 can accelerate the development of tumors in the AOM-induced CRC model. To explore the underlying mechanisms, we performed global proteomic and phosphoproteomic analysis of mice colon tissue chronically exposed to *SGG* UCN34 for 12 weeks. We compared these results to mice colonized with *S. gallolyticus* subsp. *macedonicus* (*SGM*) considered as the closest non-pathogenic relative. Colonization with *SGG* UCN34 altered the expression of 164 proteins and 725 phosphosites in colon tissue devoid of tumors as compared to the equivalent colon tissue colonized with *SGM*. The vast majority of altered proteins were associated with cancer disease. Bioinformatic analyses using IPA and ROMA software revealed significant up-regulation of multiple signaling pathways previously linked to cancer cells and their stromal microenvironment, including the three major MAPKs (ERK, JNK, p38), the PI3K/AKT/mTOR and integrin/ILK/actin cytoskeleton. These data suggest that *SGG* UCN34 affects multiple proteins in these pathways altering growth factors signaling cascades such as EGF and FGF receptors, ubiquitously expressed, and PDGFA receptors, which are mostly expressed in the colonic stromal compartment. In agreement with these results, RPPA analysis of human colon tumors enriched for *SGG* revealed up-regulation of PI3K/Akt/mTOR and MAPK pathways. A working model summarizing our results is depicted in **Fig. 6.** We hypothesize that this complex activation of multiple signaling pathways in cancer cells and their stromal microenvironment by *SGG* UCN34 can lead to cell proliferation and transformation, thus accelerating tumor development.

**Figure 6.**
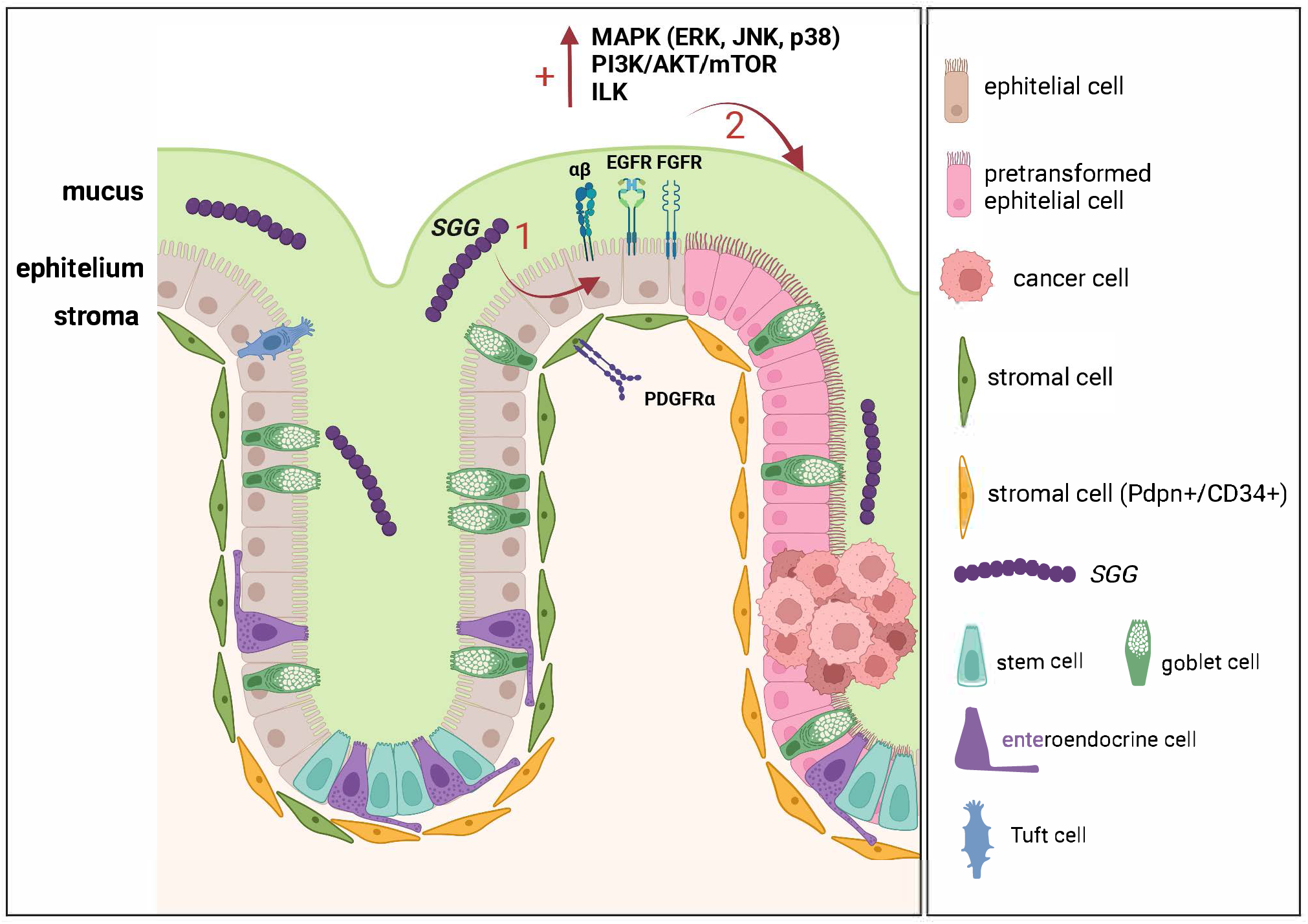
Working model of the oncogenic potential of *SGG* UCN34. Chronic colonization of host colon with *SGG* UCN34 induces the activation of MAPK cascades, PI3K/AKT/mTOR signaling and integrin/ILK/actin cytoskeleton pathways. *SGG* UCN34 alters multiple signaling pathways found downstream of EGF and FGF receptors present on colonic epithelial cells, but also downstream of PDGFA receptor present in stromal cells. *SGG* UCN34 induces the expansion of Pdpn^+^CD34^+^ cell in the stromal compartment, which regulates epithelial cells differentiation/proliferation. All these events contribute to pre-cancerous transformation of colon epithelium and acceleration of tumor development. Created with BioRender.com.

We also showed that *SGG* UCN34 contributes to early pre-cancerous transformation of murine colon epithelium, with expansion of Pdpn^+^CD34^+^ stromal niche, which have been reported to regulate epithelial cells differentiation/proliferation by producing BMP antagonists such as *Grem1* [39]. Furthermore, using an *ex vivo* organoid model, we found that *SGG* UCN34 induces the formation of abnormal organoids with compact morphology. Interestingly, this compact morphology was previously observed in human colon tumor-derived organoids [42,43]. Staining with phalloidin revealed strong disorganization of the actin cytoskeleton. These *SGG*-induced cytoskeleton rearrangements may be due to changes in the integrin/ILK/actin cytoskeleton signaling pathways [44–47].

It is worth noting that *SGG* UCN34 did not significantly increase the number and/or size of adenomas in the APC^Min/+^ mice, mirroring our previous results in the Notch/APC mice (Aymeric et al., 2017). These results indicate that the oncogenic effect of *SGG* UCN34 is influenced by host genetic factors and/or the niche environment. *SGG* UCN34 is rather an opportunistic pathobiont profiting from a favorable ecological niche (e.g., AOM-induced tumoral environment) and exerting its pro-oncogenic effect only under specific conditions. Consistent with this idea, we did not observe any direct ‘oncogenic’ cellular changes (cell proliferation, cytoskeleton reorganization, cell migration, or induction of double strand DNA breaks) *in vitro* induced by *SGG* UCN34 using short-term infection of colonic cells (**Fig. S8**). Altogether, our results indicate that *SGG* UCN34-induced alterations of colon epithelium *in vivo* are probably not direct and potentially occur through other components of this complex system, e.g., gut microbiota, mucus, stroma, immune cells and/or enteric nervous system. Future studies should focus on more complex and integrated systems such as “gut-on-a-chip” [48] to unveil the various molecular pathways underlying acceleration of tumorigenesis by *SGG* UCN34.

In parallel, the sequencing of several other *SGG* clinical isolates associated with CRC revealed their highly diverse genetic content (**Fig. S9**). This result renders the existence of a single toxin-encoded genetic island directly driving colon tumor development unlikely, differing from other CRC-associated bacteria such as pks+ *E. coli* or bft+ *B. fragilis* [49–51]. We thus hypothesize that the oncogenic potential of *SGG* may differ from one isolate to another. Genetically and geographically distant *SGG* isolates, namely *SGG* UCN34 (France) and *SGG* TX20005 (USA) accelerate the development of tumors in the AOM-induced CRC model at similar levels, but likely through different mechanisms. Indeed, *SGG* TX20005 was shown to accelerate cell proliferation *in vitro* and tumor growth *in vivo* though activation of the β-catenin pathway [23,24] which is not the case for *SGG* UCN34 (**Fig. S8A-B**). Interestingly, the genetic island encoding the type VII secretion system of *SGG* TX20005, shown to contribute to tumor development, exhibits significant differences in term of organization, expression, and secreted effectors with the majority of other *SGG* isolates exemplified by *SGG* UCN34 [52].

Finally, our group and others also previously showed that *SGG* UCN34 benefits from tumor metabolites [21] and is able to colonize the host colon in tumoral conditions by outcompeting phylogenetically-related members of the gut microbiota [22]. Building on past and present data, together with our recent paper showing significant enrichment of *SGG* in the stools of patients with adenocarcinoma [19], we propose that all *SGG* isolates have the capacity to take advantage of tumoral conditions in the host colon to multiply. In contrast, *SGG’s* ability to drive colon tumorigenesis appears strain-specific and will occur at different levels through different mechanisms. Understanding how *SGG* and other CRC-associated pathobionts increase the risk of developing colonic tumors is key to improving diagnosis and treatment of colorectal cancer.

## Material and Methods

### Bacterial strains and culture conditions

*S. gallolyticus SGG* UCN34, *SGG* TX2005 and *SGM* CIP105683T were grown at 37°C in Todd Hewitt Yeast (THY) broth in standing filled flasks or on THY agar (Difco Laboratories). Starter cultures before mice oral gavage were prepared by growing strains overnight in 50 ml THY broth. Fresh THY broth was then inoculated with the overnight culture at 1:20 ratio. Exponentially growing bacteria were harvested at 0.5 OD_600_ for the mouse gavage.

### AOM-induced CRC model in A/J mice

A/J mice were first imported from Jackson Laboratory (USA) and then bred at the Pasteur Institute animal breeding facility in SOPF condition. Eight-week-old female A/J mice were first treated with AOM (Azoxymethane; A5486; Sigma, France) at a dose of 8 mg/kg by intraperitoneal (i.p.) injection once a week for 4 weeks. After a 2 week-break, mice were treated during 3 days with a broad-spectrum antibiotic mixture including vancomycin (50 μg/g), neomycin (100 μg/g), metronidazole (100 μg/g), and amphotericin B (1 μg/g). Antibiotics were administrated by oral gavage. Additionally, mice were given ampicillin (1g/L) in drinking water for one week and then switched to antibiotic-free water 24 h prior to bacterial inoculation. Mice were orally inoculated with PBS 1X (NT), *SGM* or *SGG* using a feeding needle (~2×10^9^ CFU in 0.2 ml of PBS 1X/mouse) at a frequency of three times per week during the first week of colonization and then once a week for another 11 weeks. Stools were recovered every 2-weeks to monitor *SGG* and *SGM* numbers in the colon during the whole experiment. After 12 weeks of bacterial colonization mice were euthanized and colons removed by cutting from the rectal to the cecal end and opened longitudinally for visual evaluation. Tumor numbers were recorded, and tumor sizes measured using a digital caliper. Tumor volumes were calculated by the modified ellipsoidal formula: V = ½ (Length × Width^2^). For each mouse, all adenomas as well as adjacent tumor-free sections (1 cm^2^) were dissected and frozen by immersion in liquid nitrogen and stored at −80°C until protein extraction. For ex vivo organoid formation, adjacent tumor-free (1 cm^2^) regions for both groups *SGM* and *SGG* UCN34 were dissected and processed immediately for crypt isolation. For further histological analysis, all tissue sections (tumors, tumor-free regions) were fixed for 24 h in 4% paraformaldehyde (PFA).

### Determination of *SGG/SGM* counts in mice stools

Bacterial colonization of murine colons by *SGG* and *SGM* was determined by colony forming units (CFU) counts. Briefly, freshly collected stools were weighted and homogenized using aPrecellys homogenizer (Bertine) for 2 x 30 seconds at a frequency of 5,000 rpm. Following serial dilutions, samples were plated on Enterococcus agar selective media for counting of *SGG/SGM* colonies, exhibiting a specific pink color on these plates as described previously [53].

### Ethics statement

Animals were housed in the Institut Pasteur animal facilities accredited by the French Ministry of Agriculture for performing experiments on live rodents. Work on animals was performed in compliance with French and European regulations on care and protection of laboratory animals (EC Directive 2010/63, French Law 2013-118, February 6th, 2013). All experiments were approved by the Ethics Committee #89 and registered under the reference dap180064. Mice were housed in groups up to 7 animals per cage on poplar chips (SAFE, D0736P00Z) and were fed with irradiated food at 25 kGy (SAFE, #150SP-25). The facility has central air conditioning equipment that maintains a constant temperature of 22 ± 2 °C. Air is renewed at least 20 times per hour in animal rooms. Light is provided with a 14:10-h light:dark cycle (06:30 to 20:30). Mice were kept in polypropylene or polycarbonate cages that comply with European regulations in terms of floor surface per animal. All cages were covered with stainless steel grids and non-woven filter caps.

### Protein extraction and preparation for mass spectrometry

All proteins from previously frozen samples (tumors and healthy parts of murine colon) were extracted using AllPrep DNA/RNA/Protein Mini Kit (Qiagen). Protein concentrations were estimated using NanoDrop A280 absorbance. About 300 μg of each sample was incubated overnight at −20°C with 80% MethOH, 0.1 M Ammonium Acetate glacial (buffer 1). Proteins were then precipitated by centrifugation at 14,000 g for 15 min at 4°C and the pellets washed twice with 100 μl of buffer 1. Samples were then dried on a SpeedVac (Thermofisher) vacuum concentrator. Proteins were resuspended in 100 μl of 8 M Urea, 200 mM ammonium bicarbonate, reduced in 100 μl of 5 mM dithiotreitol (DTT) at 57°C for one hour and then alkylated with 20 μl of 55 mM iodoacetamide for 30 min at room temperature in the dark. The samples were then diluted in 200 mM ammonium bicarbonate to reach a final concentration of 1 M urea. Trypsin/LysC (Promega) was added twice at 1:100 (wt:wt) enzyme:substrate, incubated at 37°C during 2 h and then overnight. Each sample were then loaded onto a homemade C18 StageTips for desalting. Peptides were eluted using 40/60 MeCN/H2O + 0.1% formic acid and 90% of the starting material was enriched using TitansphereTM Phos-TiO kit centrifuge columns (GL Sciences, 5010-21312) as described by the manufacturer. After elution from the Spin tips, the phospho-peptides and the remaining 10% eluted peptides were vacuum concentrated to dryness and reconstituted in 0.3% trifluoroacetic acid (TFA) prior to LC-MS/MS phosphoproteome and proteome analyses.

### Mass spectrometry

Peptides for proteome analyses were separated by reverse phase liquid chromatography (LC) on an RSLCnano system (Ultimate 3000, Thermo Scientific) coupled online to an Orbitrap Exploris 480 mass spectrometer (Thermo Scientific). Peptides were trapped on a C18 column (75 μm inner diameter × 2 cm; NanoViper Acclaim PepMapTM 100, Thermo Scientific) with buffer A (2/98 MeCN/H2O in 0.1% formic acid) at a flow rate of 2.5 μL/min over 4 min. Separation was performed on a 50 cm x 75 μm C18 column (NanoViper Acclaim PepMapTM RSLC, 2 μm, 100Å, Thermo Scientific) regulated to a temperature of 50°C with a linear gradient of 2% to 30% buffer B (100% MeCN in 0.1% formic acid) at a flow rate of 300 nL/min over 220 min. MS full scans were performed in the ultrahigh-field Orbitrap mass analyzer in ranges m/z 375–1500 with a resolution of 120 000 (at m/z 200). The top 25 most intense ions were isolated and fragmented via high energy collision dissociation (HCD) activation and a resolution of 15 000 with the AGC target set to 100%. We selected ions with charge state from 2+ to 6+ for screening. Normalized collision energy (NCE) was set at 30 and the dynamic exclusion to 40 s.

For phosphoproteome analyses, LC was performed with an RSLCnano system (Ultimate 3000, Thermo Scientific) coupled online to an Orbitrap Fusion mass spectrometer (Thermo Scientific). Peptides were trapped on a C18 column (75 μm inner diameter × 2 cm; NanoViper Acclaim PepMapTM 100, Thermo Scientific) with buffer A (2/98 MeCN/H2O in 0.1% formic acid) at a flow rate of 3 μL/min over 4 min. Separation was performed on a 25 cm x 75 μm C18 column (Aurora Series, AUR2-25075C18A, 1.6 μm, C18, Ionopticks) regulated to a temperature of 55°C with a linear gradient of 2% to 34% buffer B (100% MeCN in 0.1% formic acid) at a flow rate of 150 nL/min over 100 min. Full-scan MS was acquired using an Orbitrap Analyzer with the resolution set to 120,000, and ions from each full scan were higher-energy C-trap dissociation (HCD) fragmented and analysed in the linear ion trap. The mass spectrometry proteomic and phosphoproteomic data were deposited in the ProteomeXchange Consortium via the PRIDE [54] partner repository with the dataset identifiers PXD038272 (; Zc74Ei5B); PXD038270 (; Ctnt4Ao0); PXD038268 (; FW8rInt) and PXD038267 (; blfbVkViU).

### Proteome and phosphoproteome analyses

For identification, the data were compared with the *Mus musculus* (UP000000589) SwissProt database and a common database of contaminants using Sequest HT through proteome discoverer (version 2.2). Enzyme specificity was set to trypsin and a maximum of two-missed cleavage sites were allowed. Carbamidomethylation of cysteines, Oxidized methionine and N-terminal acetylation were set as variable modifications for proteomes. Phospho-serines, - threonines and -tyrosines were also set as variable modifications in phosphoproteome analyses. Maximum allowed mass deviation was set to 10 ppm for monoisotopic precursor ions and 0.02 Da for MS/MS peaks or 0.6 Da for phosphoproteome analyses. FDR calculation used Percolator [55] and was set to 1% at the peptide level for the whole study. The resulting files were further processed using myProMS [56] v3.9.3 (https://github.com/bioinfo-pf-curie/myproms).

Label free quantification was performed by peptide Extracted Ion Chromatograms (XICs), reextracted across all conditions (healthy/tumor-free, tumor, polyp and mix samples, see PXD038272, PXD038270, PXD038268 and PXD038267) and computed with MassChroQ version 2.2.1 [57]. Polyp samples were not used in this study since histological analysis demonstrated that they correspond to lymphoid aggregates (LA, Fig. S1E). For differential analyses, XICs from proteotypic peptides (shared between compared conditions for proteomes, no matching constraints for phosphoproteomes) with at most two-missed cleavages were used. Replicate *SGM* m8-H which had more than 76% missing values was excluded from the proteome differential analyses. Median and scale normalization was applied on the total signal to correct the XICs of each biological replicate for injection and global variance biases. To estimate the significance of the change in protein abundance, a linear model (adjusted on peptides and biological replicates) was performed, and *p*-values were adjusted with a Benjamini–Hochberg FDR procedure. For proteome analyses, proteins were considered in the analysis only when they were found with at least 5 total peptides. Then, proteins with an adjusted *p*-value ≤ 0.05 were considered significantly enriched in sample comparison. For phosphoproteome analyses, phosphosites were analyzed individually. The threshold was also 5 total phosphopeptides to consider a phosphosite for the downstream analysis. Then, phosphosites with an adjusted *p*-value ≤ 0.05 were considered significantly changed in sample comparisons. Unique phosphosites (only detected in one of the groups in each comparison) were also included when identified in at least 5 biological replicates. In addition, we considered the extent of the change in expression to be significant if higher than or equal to 2 for up-regulated entities or lower or equal to 0.5 for down-regulated entities. For the other bioinformatic analyses, label-free quantification (LFQ) was performed following the algorithm as described [58] for each sample after peptide XIC normalization as described above. The resulting LFQ intensities were used as protein (all peptide ≥ 3) or phosphosite (all peptides ≥ 2) abundance. For PCA and ROMA analyses, datasets were further filtered to remove entities with more than 34% of missing values across all samples used. The LFQ values were log10-transmformed and the remaining missing values (around 5% for proteome and 15% for phosphoproteome) were imputed using the R package missMDA [59] to produce complete matrices.

### Quantification of pathway activity with ROMA

ROMA (Representation and quantification Of Module Activities) is a gene-set-based quantification algorithm [31] with the MSigDB C2 Canonical pathways sub-collection as pathway database composed of 2232 pathways from Reactome (http://www.reactome.org), KEGG (http://www.pathway.jp), Pathway Interaction Database (http://pid.nci.nih.gov), BioCarta (http://cgap.nci.nih.gov/Pathways/BioCarta_Pathways) and others. ROMA was used to investigate potential changes in pathway activity between the 3 groups (*SGG* UCN34 tumor-free, *SGM* tumor-free and *SGG* UCN34 tumor) based on phosphoproteomic data. In this study, we used an R implementation of ROMA available at https://github.com/Albluca/rRoma. Only the 16 most representative samples of the 3 groups were used for the analysis (samples *SGM* healthy m6, *SGM* healthy m9, *SGG* healthy m10 and *SGG* tumor m14 were removed due to their outlier behavior as illustrated in Fig. 2C). A data reduction to a single phosphosite per protein/gene was performed to switch from a site-centric to a gene-centric dataset. This reduction was achieved with an ANOVA across groups and selection of the site with the lowest *p*-value (“most informative” site) when multiple sites were quantified for the same protein. 2784 of the 3188 mouse genes implicated were then converted into their human counterparts with the Biomart resource (https://m.ensembl.org/biomart, database version 100) prior to ROMA analysis against the MSigDB (http://www.gsea-msigdb.org) human C2 Canonical pathways sub-collection (version 7.1) as pathway database. 1752 genes from the dataset matched 666 of the 2232 pathways.

### Ingenuity Pathway Analysis (IPA)

We used IPA (QIAGEN, http://www.ingenuity.com/) to characterize two sets of proteins: (i) one corresponding to the list of proteome changes between the tumor-free *SGG* UCN34 group and the tumor-free *SGM* group (164 proteins of which 163 IDs were found in the IPA database) and (ii) and one to the list of phosphoproteome changes between the tumor-free *SGG* UCN34 group and the tumor-free *SGM* group (597 proteins of which all IDs were found in the IPA database). For the list of phosphoproteins, we first removed phosphosites that were found in conjunction with other phosphosites on the same protein, and in that case, we retained only the site with the lowest *p*-value (in total 725 phosphosites mapped to 597 proteins). Lists containing differentially expressed proteins with their corresponding log2 ratio of expression values for both data sets (proteome and phosphoproteome) were uploaded and analyzed individually. We only considered proteins and phosphoproteins measurements where we had at least 5 non-zero expression measurements in at least one of the conditions. For the cases where we had no expression in one of the conditions, leading to non-finite log2 ratios, we set the log2 ratio values to be equal to +10 [resp. −10] in the case of an upregulation [resp. down-regulation] in the tumor-free *SGG* UCN34 group. This value corresponded to the upper end (in absolute value) of log2 ratios for comparisons of measurements containing no zero-values. This measurement (log2 ratio) was used by IPA to calculate directionality (z-scores) in the analysis and is displayed in color on pathways and networks (red for up-regulated proteins and green for down-regulated proteins). We used the “Core Analysis” function to relate the expression to known diseases, biological functions, and canonical pathways. This software is based on computer algorithms that analyze the functional connectivity of genes from information obtained from the IPA database. Biological Functions with a corrected *p*-value < 0.05 (Fisher’s exact test) were considered to be statistically significant. The analysis of canonical pathways identified the pathways from the IPA library of canonical pathways which were most relevant to the input data set, based on a test of enrichment (Fisher’s exact test).

### Crypt Isolation and Organoid Formation

Half centimeter sections of tumor-free colonic tissue were removed and washed in cold PBS 1X. The tissue was further fragmented in smaller pieces using a scalpel and washed in fresh cold PBS 1X by pipetting 5-times with pre-cut P1000 tips. After sedimentation of the pieces, the supernatant was removed and the fragments were digested in HBSS with calcium and magnesium, 2.5% fetal bovine serum (FBS), 100U/ml penicillin-streptomycin, 1mg/ml collagenase from Clostridium histolyticum (Sigma C2139), and 1μM ROCK1 inhibitor (Y-27632, StemCell technologies), for 60 min in a thermomixer at 37°C with 800 rpm agitation. After a quick spin, the fragments were further digested using Tryple express (Gibco) with 1μM ROCK1 inhibitor, for 20 min at 37°C with 800rpm agitation. The samples were washed with PBS 1X, left to sediment and the supernatant was collected, centifuged and the pellet resuspended with Matrigel Growth Factor Reduced Basemen Membrane Matrix (Corning) in order to make 2 drops of 30 μl each in a 48-well plate. After 20 min at 37°C, the Matrigel drops were overlayed with 300 μl of Medium containing Advanced DMEM, P/S, N2 and B27 (all from Gibco), lacking Noggin, R-spondin, Wnt3a, and EGF. The observation of organoids formation was done at day 6.

### Immunofluorescence

Tissues and organoids were fixed with 4% paraformaldehyde (Electron Microscopy Science) overnight at 4°C. Tissues were embedded in OCT compound (Fisher Scientific) and stored at −80°C. Frozen blocks were cut at 10 μm thickness, and sections were collected onto Superfrost Plus slides (VWR international). Tissue sections and organoids were permeabilized with 0.5% triton X-100 (Sigma Aldrich) in PBS 1X at RT for 40 min and incubated with a blocking buffer containing 3% bovine serum albumin (BSA, Sigma Aldrich) in PBS 1X RT for 40 min. Samples were incubated overnight at 4°C with primary antibodies diluted in 0.1% triton X-100, 1% BSA in PBS 1X, washed, incubated with secondary antibodies for 1 h at room temperature, washed, and stained for 20 min with 1 μg /ml DAPI (Invitrogen). Finally, slides were mounted with ProLong™ gold antifade mounting medium (ThermoFisher). The following antibodies were used: anti-CD34 coupled with eF660 (clone RAM34) (eBioscience), anti-E-cadherin (clone ECCD-2) (Takara), goat anti-rat488 (Thermo Fisher), anti-Pdpn (gift from A. Farr, University of Washington, Seattle), and goat-anti-hamster 546 (Thermo Fisher), anti-Ki67 coupled with eF660 (clone SolA15) (eBioscience), phalloidin coupled with AF488 (Thermo Fisher).

### Histology and immunohistochemistry

Colon fragments fixed for 48 hours in 10% neutral-buffered formalin were embedded in paraffin. Four-μm-thick sections were cut and stained with hematoxylin and eosin (H&E) staining. Histological evaluation was performed by an histopathologist in a blind fashion. Characterization of LA was done by immunostaining performed on Leica Bond RX using anti-CD3 (Dako, Ref. A0452) antibody, and hematoxylin staining. Slides were then scanned using Axioscan Z1 Zeiss slide scanner and images were analyzed with the Zen 2.6 software.

### Patient samples

Colon biopsy samples were collected from patients at the University of Texas M. D. Anderson Cancer Center (MDACC), Houston, Texas. The cohort contains mostly stage II and III tumors. The patients had previously given written informed consent for use of their samples in future colorectal cancer research. Patient identifiers were anonymized. Collection and handling of patient samples were carried out in strict accordance to protocols approved by the institutional review board at MDACC and Texas A&M Health Science Center.

Detection of *SGG* in the colon biopsy samples was described previously (Kumar et al. 2017).

### Reverse phase protein array (RPPA)

This was performed in the RPPA core at MDACC using established protocols. Briefly, frozen tumors were lysed and protein extracted. Lysates were serially diluted in 5 two-fold dilutions and printed on nitrocellulose-coated slides using an Aushon Biosystems 2470 arrayer. Slides were probed with primary antibodies selected to represent the breadth of cell signaling and repair pathways [60] conditioned on a strict validation process as previously described [61], followed by detection with appropriate biotinylated secondary antibodies and streptavidin-conjugated horseradish peroxidase (HRP). The slides were analyzed using Array-Pro Analyzer software (MediaCybernetics) to generate spot intensity, which was adjusted using “control spots” to correct spatial bias [62]. A fitted curve (“Supercurve”) was created for each protein using a non-parametric, monotone increasing B-spline model [34]. Relative protein levels were estimated using SuperCurve GUI [63]. Slide quality was assessed using a QC metric [64] and only slides greater than 0.8 on a 0-1 scale were included for further processing. Protein measurements were corrected for loading as described [63,65] using bidirectional median centering across samples and antibodies.

### Statistical Analysis

Mann-Whitney nonparametric test was used to test for statistical significance of the differences between the different group parameters. *p* values of less than 0.05 were considered statistically significant.

## Acknowledgements

This work was supported by the Institut National contre le Cancer (INCA, grant PLBIO16-025) and from the French Government’s Investissement d’Avenir program, Laboratoire d’Excellence Integrative Biology of Emerging Infectious Diseases (grant no. ANR-10-LABX-62-IBEID). EPK received a 2-year Roux-Cantarini post-doctoral fellowship.

We are grateful to Johan Bedel and Magali Tichit of the Institut Pasteur Histological Platform for paraffin embedding, sectioning, histochemistry and immunohistochemistry, Karim Sebastien and Marion Bérard for technical help with animal experimentation and all the zootechnicians from the Animal Breeding Facility from Pasteur Institute.

We thank Dmitry Ershov for image analysis for DNA damage (yH2AX) quantifications, Adriana Mihalache, an histopathologist for tumor grades scoring, Bruno Périchon and Thomas Cokelaer for *SGG* isolate sequencing, Laurence du Merle for technical assistance with bacterial cultures and CFU counts as well as Valentin Sabatet from the Institut Curie Proteomics Platform for protein quantification and Olivier Mirabeau for bioinformatic help.

## Author’s contributions

P-K.E. and D.S. conceived the study; P-K.E., N.G., D.F., S.K. and D.J. conducted the experiments; P-K.E., N.G., X.Y., L.D., P.L. and P.P. analyzed the data; L.D. and D.F. performed the proteomics and phosphoproteomics analyses; P.P. performed bioinformatic analyses; R-M.C., S.P., T-C.P., X.Y. and D.S. participated in funding acquisition. E.P-K. and S.D., wrote the paper. All authors edited the manuscript and provided the critical advice.

