## Supplemental Figures 1 to 7 for "The oncogenic role of *Streptococcus gallolyticus subsp*. *gallolyticus* is linked to activation of multiple cancer-related signaling pathways"

### Supplementary figure legends

**Figure S1. Two different strains of *SGG*, UCN34 and TX20005, induce development of similar numbers of colonic tumors in AOM-induced CRC model. Most of the observed tumors for *SGG* UCN34 and TX20005 were low-grade adenomas. A.** Histological (H&E) sections of several examples of colon tumors found in the AOM-induced CRC mouse model for each experimental group: NT, *SGM*, *SGG* UCN34, and *SGG* TX20005. **B.** Graph presents the sum of tumor numbers per mouse. Macroscopic tumors were evaluated by an experimented observer. ns, not significant; *, p < 0.05; **, p < 0.01; Mann-Whitney test.

**Figure S2. *SGG* UCN34 does not induce significant acceleration of tumorigenesis in APC$^{min-/+}$ CRC model.** The effect of SGG UCN34 vs *SGM* on the development of adenomas in small intestine was examined in APC$^{min-/+}$ CRC model, using the experimental design as shown in **A**. **B**. The sum of adenomas counted from the whole small intestine for each mouse per group and the sum of adenomas volume counted from the whole small intestine for each mouse per group. ns, not significant; *, $p < 0.05$; **, p < 0.01; Mann-Whitney test. **C**. Tissue sections with fecal material were collected from the small intestine and colon, homogenized, and serial dilutions were plated onto Enterococcus Selective Agar plates to count *SGG* UCN34 cells.

**Figure S3. Canonical pathways altered by *SGG* UCN34 visualized using IPA software.** The data set used in this analysis was phosphoproteins differentially expressed between *SGG* UCN34 and *SGM* detected in macroscopically tumor-free colonic tissue. Canonical pathways that were most significant to the data set were identified from the QIAGEN Ingenuity Pathway Analysis library of canonical pathways. Canonical pathways with *p*-values < 0.05 (Fischer's exact test) were considered to be statistically significant. The activation Z-score was calculated to predict activation or inhibition of transcriptional regulators based on published findings accessible through the Ingenuity knowledge base. Regulators with Z-score greater than 2 (positive Z-score) or less than −2 (negative Z-score) were considered to be significantly activated (orange) or inhibited (blue). Regulators with Z-score of 0 are represented in white and those for which the Z-score couldn't be calculated are shown in grey.

**Figure S4. IPA identified the ERK/MAPK signaling (A) and mTOR signaling (B) signaling pathways as enriched in colon tissue colonized by *SGG* UCN34 vs *SGM*.** The data set used in this analysis was phosphoproteins differentially expressed between *SGG* UCN34 and *SGM* detected in macroscopically tumor-free colonic tissue. Nodes represent molecules in a pathway, while the biological relationship between nodes is represented by a line (edge). Edges are supported by at least one reference in the Ingenuity Knowledge Base. The intensity of color in a node indicates the degree of up- (red) or down- (green) regulation. Nodes that are red and green represent the increased and decreased measurements respectively. Nodes in orange represents predicted (hypothetical) activation and nodes in blue predicted (hypothetical) inhibition. Nodes are displayed using shapes that represent the functional class of a gene product (Circle = Other, Nested Circle = Group or Complex, Rhombus = Peptidase, Square = Cytokine, Triangle = Kinase, Vertical ellipse = Transmembrane receptor). Edges are marked with symbols to represent the relationship between nodes (Line only = Binding only, Flat line = inhibits, Solid arrow = Acts on, Solid arrow with flat line = inhibits and acts on, Open circle = leads to, Open arrow = translocates to). An orange line indicates predicted upregulation, whereas a blue line indicates predicted downregulation. A yellow line indicates expression being contradictory to the prediction. Gray line indicates that direction of change is not predicted. Solid or broken edges indicate direct or indirect relationships, respectively.

**Figure S5. IPA identified the actin cytoskeleton signaling (A), ILK signaling (B), and integrin signaling pathway (C) as enriched in colon tissue colonized by *SGG* UCN34 vs *SGM*.** The data set used in this analysis was phosphoproteins differentially expressed between *SGG* UCN34 and *SGM* detected in macroscopically tumor-free colonic tissue. Nodes represent molecules in a pathway, while the biological relationship between nodes is represented by a line (edge). Edges are supported by at least one reference in the Ingenuity Knowledge Base. The intensity of color in a node indicates the degree of up- (red) or down- (green) regulation. Nodes that are red and green represent the increased and decreased measurements respectively. Nodes in orange represents predicted (hypothetical) activation and nodes in blue predicted (hypothetical) inhibition. Nodes are displayed using shapes that represent the functional class of a gene product (Circle = Other, Nested Circle = Group or Complex, Rhombus = Peptidase, Square = Cytokine, Triangle = Kinase, Vertical ellipse = Transmembrane receptor). Edges are marked with symbols to represent the relationship between nodes (Line only = Binding only, Flat line = inhibits, Solid arrow = Acts on, Solid arrow with flat line = inhibits and acts on, Open circle = leads to, Open arrow = translocate to). An orange line indicates predicted upregulation, whereas a blue line indicates predicted downregulation. A yellow line indicates expression being contradictory to the prediction. Gray line indicates that the direction of change is not predicted. Solid or broken edges indicate direct or indirect relationships, respectively.

**Figure S6. Expression of major receptors and stromal markers in human Colorectal Cancer.** Expression of selected genes and tSNE visualization of epithelial cells (**A**) and stromal cells (**B**) from single cells RNASeq dataset of human colorectal cancer and adjacent normal tissue, published by Pelka et al., Cell 2021[1]. Expression of *Pdgfra* is mostly detected in *Pdpn*$^+$ stromal cells, which include *Cd34*$^+$ subsets producing *Grem1*. Expression of *Cd34* is also detected in *Vwf*$^+$ endothelial cells (**C**).

**Figure S7. Sample photos of organoids with cystic/normal and compact morphology.** Colored confocal pictures show DAPI nuclei labeling (blue), Phalloidin (green) and KI67/proliferation marker (red). Scale bar: 50 µm.

**Figure S8. *SGG* UCN34 does not induce cell proliferation, β-catenin activation, cell migration, cytoskeletal rearrangements, and DNA damage. A**. Cell proliferation assays. HT29 and HCT116 cells were co-cultured with SGG UCN34, SGM or media only for 24 hours and viable cell numbers enumerated. **B**. The level of β-catenin was determined by Western blot assays using total cell lysates from cells co-cultured with *SGG* UCN34, CIP 105428T (isolated from Koala's feces) or *SGM* or media only (NT). **C**. Transwell cell migration assay was performed over 16h after 24h of A549 cells co-culture with SGG UCN34 or SGM. **D**. Confocal microscope images (63X) of Caco-2 cells infected with SGG UCN34 or control SGM during 24h at an MOI of 1. Upper Panel: Pictures show DAPI nuclei labeling (blue), phalloidin/actin (red) and anti-SGG or anti-SGM (green); Lower Panel: Pictures show DAPI nuclei labeling (blue), phalloidin/actin (red) and E-cadherin (green) or occluding (green). Scale bar = 5 µm. **E.** Graph presenting the number of yH2AX positive cells (≥ 3 yH2AX foci/cell) in human normal colon cell line FHC incubated for 24h with SGM or SGG UCN34, for 4h with *E. coli* pks+ or irradiated at 5Gy and incubated 1h after irradiation. Cells were fixed and stained with yH2AX (marker of DNA damage foci) and DAPI and imaged by confocal microscopy at 63X objective. The determination of yH2AX positive cells was done using automated imaging analysis software Icy.

**Figure S9.** Phylogenetic tree of the SBSEC complex showing that *SGM* CIP105683T is genetically closely related to *SGG* UCN34

**Supplemental Table S1.** PROTEOME: List of 164 proteins differencially detected between tumor-free colon colonized by SGG UCN34 and SGM out of 7241 identified proteins in total.

**Supplemental Table S2.** PHOSPHOPROTEOME: List of 725 phosphosites/598 proteins differencially detected between tumor-free colon colonized by SGG UCN34 and SGM out of 12005 phosphosites/4102 proteins identified in total.

**Supplemental Table S3.** List of pathways up and down-regulated based on phosphoproteome changes between 3 groups: tumor-free SGG UCN34, tumor-free SGM and tumor SGG using ROMA analysis tool.

**Supplemental Table S4.** RPPA analysis on human colon tumors enriched with SGG vs negative ones.

**Supplemental Table S5.** List of phosphoproteins from all up-regulated pathways detected by ROMA.

### Supplementary Material & Methods

##### $APC^{Min/+}$ mouse CRC model

$APC^{Min/+}$ mice were provided by the Institut Pasteur animal breeding facility. Five-week-old female $APC^{Min/+}$ mice [2] were first treated with a broad-spectrum antibiotic cocktail including vancomycin (50 µg per g), neomycin (100 µg per g), metronidazole (100 µg per g), amphotericin B (1 µg per g) and ampicillin (1g per L) for 8 days, as previously described (Reikvam et al. 2011) and switched to antibiotic-free water 24 hours prior to bacterial inoculation. Oral gavage of mice with *SGM* or *SGG* was done using a feeding needle (~$2x10^9$ cfu in 0.2ml of PBS/mouse) at a frequency of three times per week during the first week of colonization and then once a week for another 12 weeks. Stools were collected every week before new bacterial inoculation to estimate the number of bacteria and ensure good colonization state. After 13 weeks of bacterial colonization mice were euthanized and small intestines and colons were removed by surgery and opened longitudinally for visual evaluation. Adenoma numbers were counted under the binocular loupe. To estimate tumor volume by external caliper, the greatest longitudinal diameter (length) and the greatest transverse diameter (width) were determined of each adenomas/mouse. Tumor volume was calculated by the modified ellipsoidal formula: $V = ½ (Length × Width^2)$.

##### Cell culture

The human normal colon epithelial cell lines, FHC (ATCC: CRL-1831) were cultured in DMEM/F12 medium (Gibco, France) supplemented with 10% heat-inactivated calf serum and additional factors (25 mM HEPES; 10 ng/mL cholera toxin; 0.005 mg/mL insulin; 0.005 mg/mL transferrin; 100 ng/mL hydrocortisone; EFG 20 ng/mL; 10% SVF) to sustain their growth and could be passed 5-10 times only. The human cancerous cell lines HT-29 (ATCC: HTB-38), HCT-116 (CCL-247), Caco2 (HTB-37) and A549 (CRM-CCL-185) were cultivated in DMEM with 10% heat-inactivated calf serum and supplemented with 25 mM HEPES. The cells were cultured in ventilated T75 flasks at 37 °C and 5 % $CO_2$.

##### Proliferation assay

Cells were seeded onto the wells of 6-well plates at $1x10^4$ cells per well and incubated for 16-20 hours. Stationary phase bacteria were scraped from fresh THY plates (o.n. culture), washed with sterile phosphate buffered saline, pH 7.4 (PBS) and resuspended in the appropriate cell culture media. Bacteria were added to the wells at 1x$10^4$ CFU/well for SGG UCN34 and at 1x$10^5$ CFU/well for SGM and incubated for 24 hours. Trimethoprim was added at 50 µg/ml final concentration after 6 hours of incubation to prevent bacterial growth leading to media acidification. To estimate cell numbers after 24h of bacteria-cell co-culture, cells were detached by trypsin treatment, stained with trypan blue and counted in a TC20 automated cell counter (Biorad).

**Western Blotting**

Cells were cultured in the appropriate medium in the presence or absence (NT) of bacteria (UCN34, multiplicity of infection (MOI)=1; CIP 105428T, MOI=1 or SGM, MOI=10) for 24 hours and washed with sterile PBS three times. Cells were lysed by scraping into Laemmli buffer (0.125M Tris-HCl; pH 6.8, 4% SDS, 20% glycerol, 2mM DTT, 1X Protease Inhibitor Cocktail) and boiling for 10 min. Cells were cultured in the appropriate medium in the presence or absence (NT) of bacteria (UCN34, MOI=1 or SGM, MOI=10) for 24 hours and washed with sterile PBS 1X three times. Cells were lysed by scraping into Laemmli buffer (0,125M Tris-HCl; ph 6,8, 4% SDS,20% glycerol, 2mM DTT, 1X Protease Inhibitor Cocktail) and boiling for 10 min. The resulting lysates were then centrifuged, and protein concentrations were estimated using NanoDrop A280 absorbance. Proteins (20µg) were separated by 4-15% Mini-PROTEAN TGX stain-free gels (Bio-Rad), transferred to PVDF membranes (Trans-Blot Turbo, Bio-Rad), blocked by incubation with 5% of milk for 1 h and hybridized overnight at 4°C with primary antibody diluted in 5% of milk. Antibodies used were purified mouse antibody against β-catenin (1:1000; BD; Ref. 610154), monoclonal mouse anti β-actin (1:10000; SIGMA; Ref. A5441). Membranes were probed with goat anti-mouse secondary antibodies conjugated to Alexa Fluor 680 (Invitrogen) or Alexa Fluor 800 (Invitrogen). Blots were imaged and quantified with the Odyssey Infrared Imaging System (LI-COR Biosciences, Lincoln) and Odyssey software.

**Transwell cell migration assay**

Cells were seeded onto the wells of 6-well plates at 1x$10^5$ cells per well and incubated for 16-20 hours. Stationary phase bacteria were scraped from fresh THY plates (o.n. culture), washed with sterile phosphate buffered saline, pH 7.4 (PBS) and resuspended in the appropriate cell culture media without fetal bovine serum. Bacteria were added to the wells at 1x$10^4$ CFU/well for SGG UCN34 and at 1x$10^5$ CFU/well for SGM and incubated for 24 hours. 300 µL of the cell suspension (50 000 cells/mL in serum-free media) for each experimental condition (SGG

UCN34, SGM) was then added to the upper migration chamber (Permeable Support for 24-well Plate with 8.0 µm Transparent PET Membrane, Corning; Ref. 353097) and 500 µL of culture media with 10% of fetal bovine serum was added to each well of the lower chamber (24-well plates). Media from upper and lower chambers were also complemented with penicillin/streptomycin (1X) to avoid further bacterial growth. The plates were incubated in a tissue culture incubator at 37°C with 5% CO2 for 24 hours. After the incubation period, the media from the upper chamber was aspirated and cells were fixed with 4% of paraformaldehyde (PFA) for 20 min. Then cells were stained with 0.1% crystal violet in 10 % ethanol for 30 min. We then performed several washes with $H_2O$ and used cotton swabs to remove the remaining non-migratory cells from the interior part of the insert. Cell migration from the upper to the lower side of the filter was observed under light microscopy. The photos were taken at objective 10X (14 different fields/chamber). Number of migratory cells were calculated using Cell Counter plugins of Fiji software.

**Immunostaining and microscopy**

HT-29-MTX cells were grown over 18-21 days on glass coverslips placed in a 24-well plate to permit full polarization. Cells were then infected with *SGG UCN34* or *SGM* at different MOI (1, 10, 1000) and for different periods of time (1h, 4h, 6h, 24h). Following infection, cells were washed 5 times with PBS 1X, fixed with 4% paraformaldehyde for 20 min and rinsed with PBS. Cells were then permeabilized using PBS 1X + 0.5% Triton X-100 for 10 min at room temperature (RT). Cells were washed again, and unspecific binding sites saturated using PBS + 2% BSA for 20min at RT. The following primary antibodies were used: rabbit anti-E-cadherin (Cell Signaling, Ref. 3195), rabbit anti-occludin (Zymed, Ref. 71-1500), rabbit anti-*SGG* or anti-*SGM* as described previously [3]. Coverslips were rinsed twice with PBS 1X and incubated with the secondary antibody: Alexa Fluor™ 647 - Phalloidin (Invitrogen), goat anti-mouse Alexa Fluor-488 (Invitrogen), goat anti-rabbit Alexa Fluor 488/633 (Invitrogen). Then, coverslips were rinsed with PBS 1X and incubated with DAPI (0.5 ug/mL in PBS 1X) for 5 minutes at RT. Coverslips were washed with PBS 1X a last time before being mounted on slides using Fluoromount-G, Invitrogen. Coverslip edges were sealed using nail polish to avoid drying. Slides were kept in the dark at 4°C. Samples were observed using a Leica TCS confocal microscope SP8, with a 63X oil immersion objective. Images were processed with the freely available Fiji software.

For experiments to detect DNA damage, FHC cells ($2x10^5$ cells/well) were first seeded into 6-well plates with glass coverslips on the bottom and allowed to attach over 16-20h. Cells were then infected with *SGG* UCN34 ($6,5x10^5$ CFU/ml) or *SGM* ($6,5x10^5$ CFU/ml) for 24h. For a positive control of DNA damage induction, we used genotoxin producing *E. coli* pks+ IHE3034 bacterial strain [4]. We followed the previously described protocol [5]. Briefly we infected FHC cells with $2,5x10^{10}$ CFU of *E. coli* pks+ for 4h. All bacteria were diluted in DMEM (Gibco, Ref. 12320032, low glucose, pyruvate, HEPES) complemented with 10% of heat inactivated FBS. For NT condition only 2 ml of fresh media was added. Another positive control for DNA damage was the FHC cells that were irradiated with the $^{137}Cs$ unit of IBL-637 (ORIS, France) at room temperature (RT) at the dose of 5Gy and fixed 1h post-irradiation. Cells were when fixed and stained using mouse primary antibodies against yH2AX (Cell Signaling, Ref. 2595) followed by a secondary antibody conjugated to a fluorescent molecule (goat anti-mouse Alexa fluor 680, Invitrogen). For *SGG* and *SGM* bacteria detection we have used polyclonal rabbit non-commercial antibodies specific to either *SGG* or *SGM* [3] followed by a secondary antibody conjugated to a fluorescent molecule (goat anti-rabbit Alexa fluor 488, Invitrogen). At least 20 fields for each condition were imaged using a 63X objective on a Leica TCS confocal microscope SP8. Image analysis was performed with a protocol in Icy [6]. Briefly: nuclei were segmented by thresholding and separated from touching objects with distance-based watershed method. Spots corresponding to damage sites were detected using the wavelet spot detector, then inclusion analysis was used to assign each damage spot to a nuclei. The resulting data was analyzed in Python using Pandas [7].

**Whole genome sequencing of *SGG* strains.**

DNA for whole-genome sequencing was isolated using the Qiagen Blood and Tissue DNA Isolation Kit (Qiagen, USA), according to the manufacturer's instructions. Quantification of extracted DNA was measured with the “Qubit 2 Fluorometer”. DNA libraries were prepared using the Illumina Nextera Kit (Illumina, USA) and sequenced on an Illumina MiSeq instrument. De novo assemblies were performed by using Sequana project [8], the Sequana denovo pipeline v0.8.5 (https://github.com/sequana/denovo) that includes assembly such as Canu software [9] and standard quality controls including remapping and coverage [10]. Alignment of whole genome sequences was performed by using MAUVE software [11]. NCBI tree was designed by using T-Rex web server [12].

### Supplementary Figures – S1

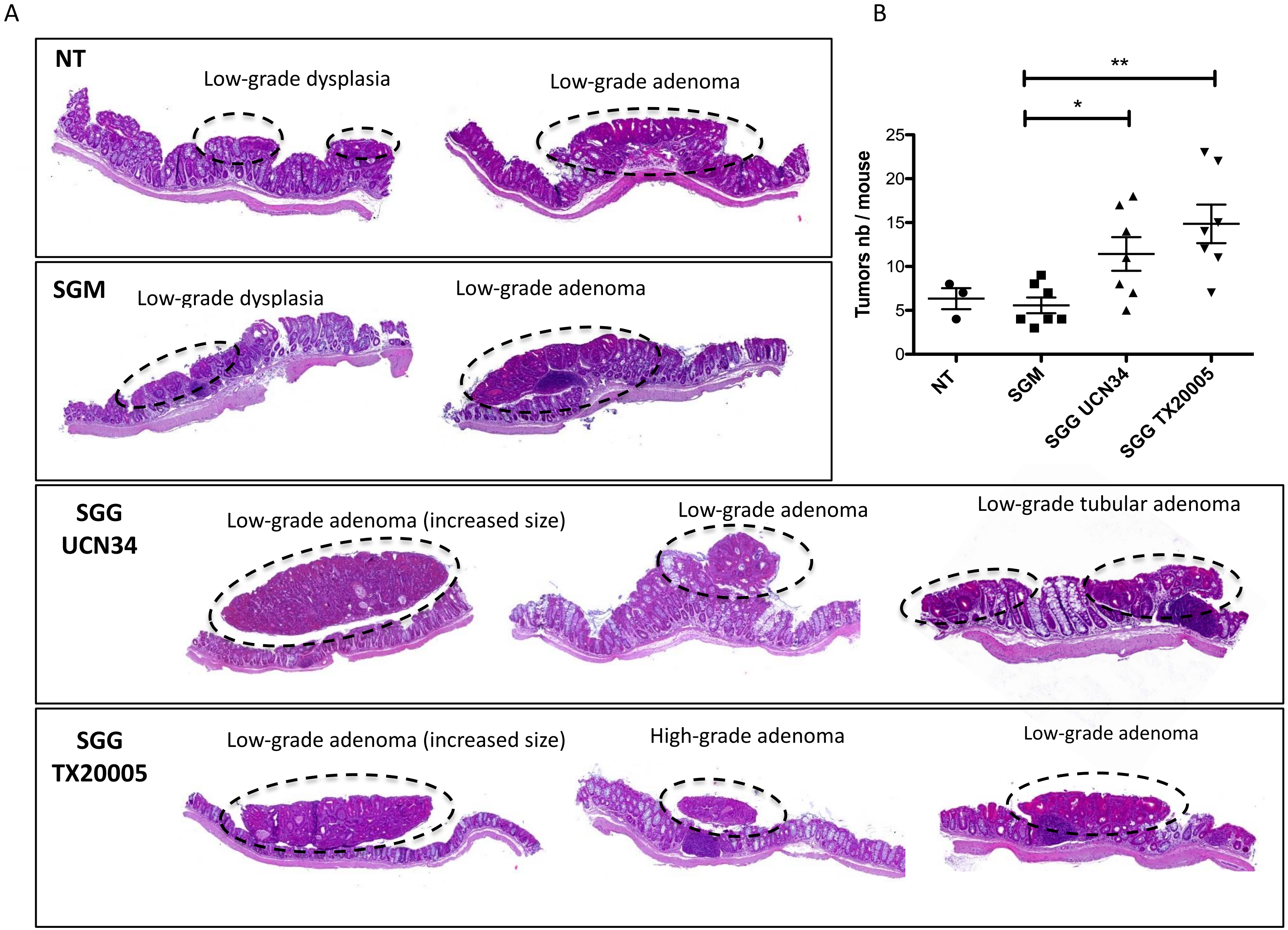

### Supplementary Figures – S2

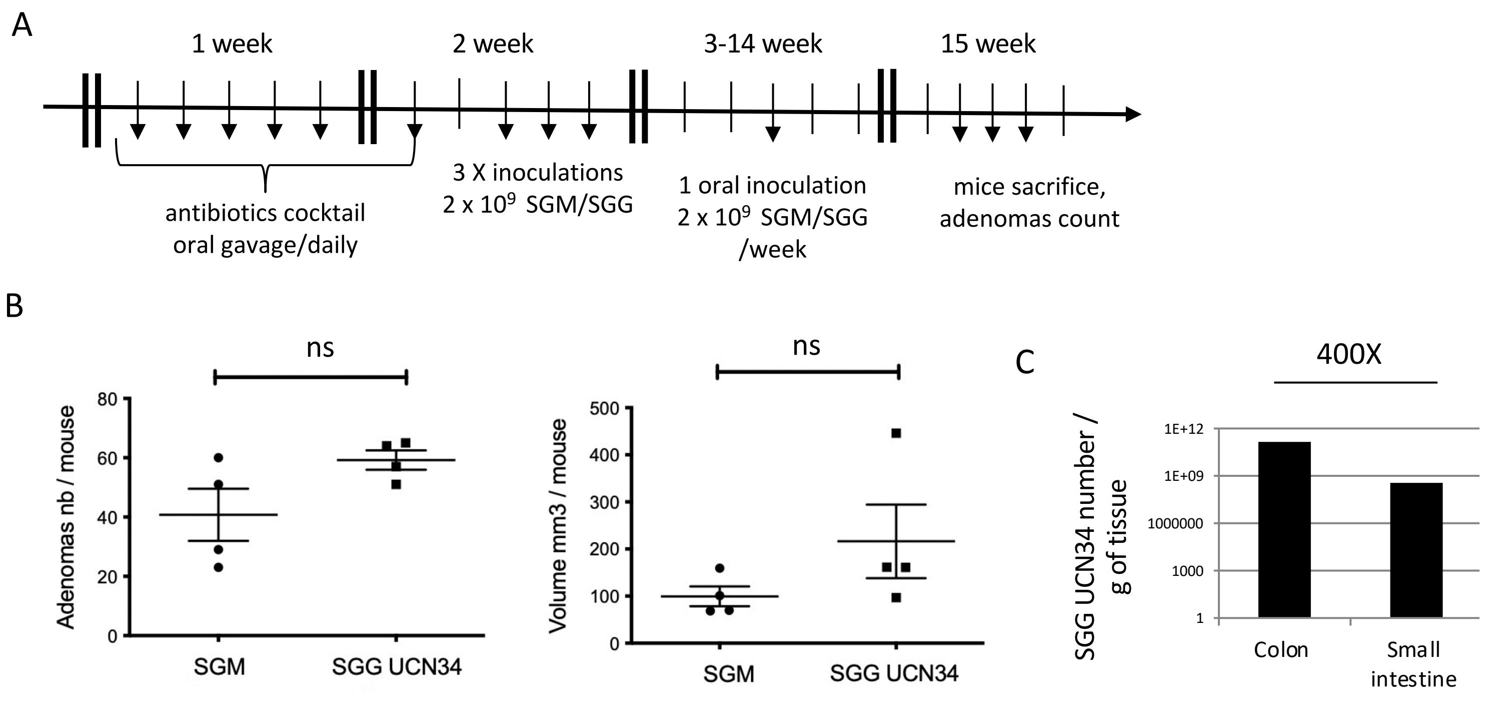

### Supplementary Figures – S3

#### IPA: PHOSPHOPROTEOME (SGG tumor-free / SGM tumor-free: total up/down : **583**)

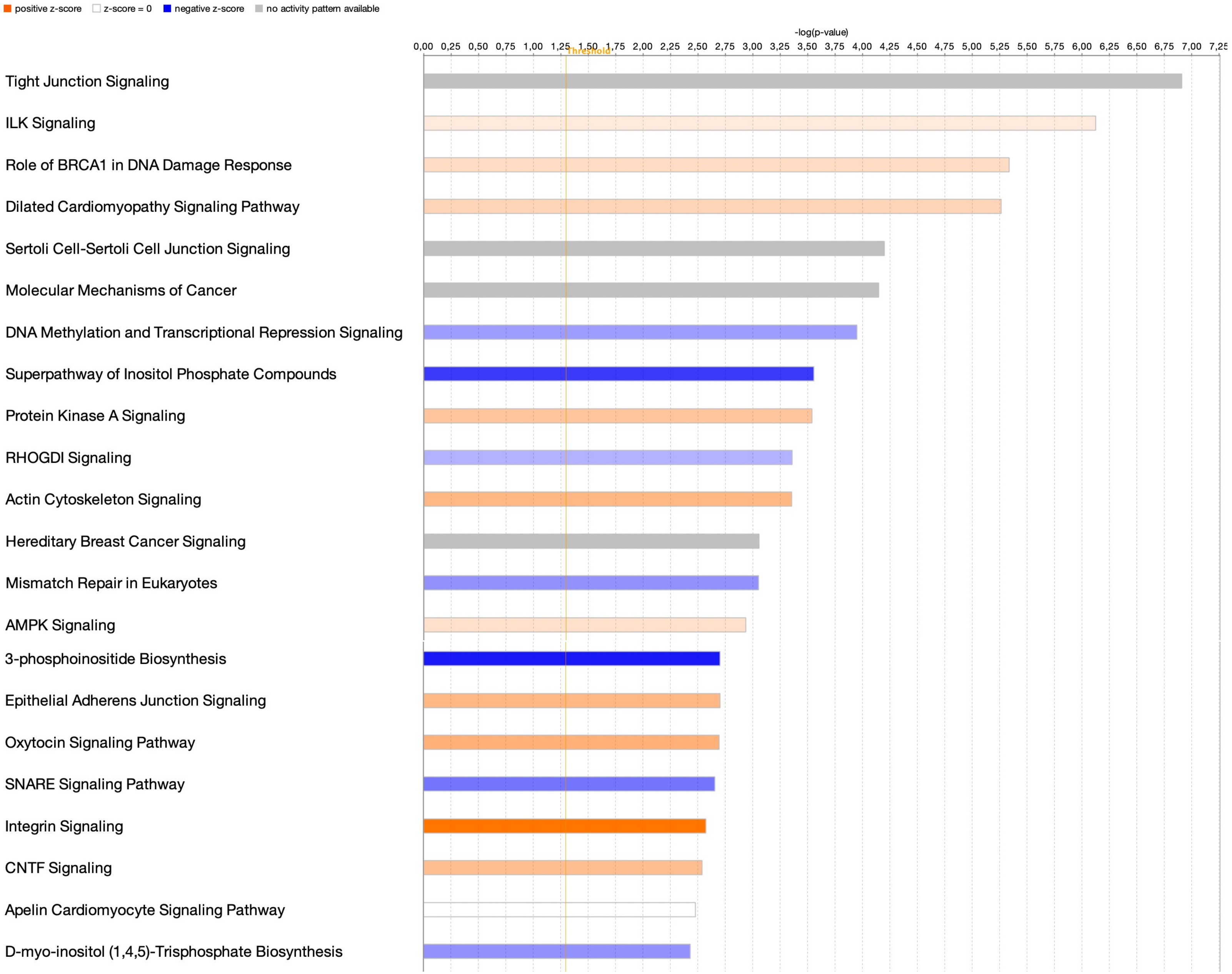

### Supplementary Figures – S4

A

#### ERK/MAPK signaling

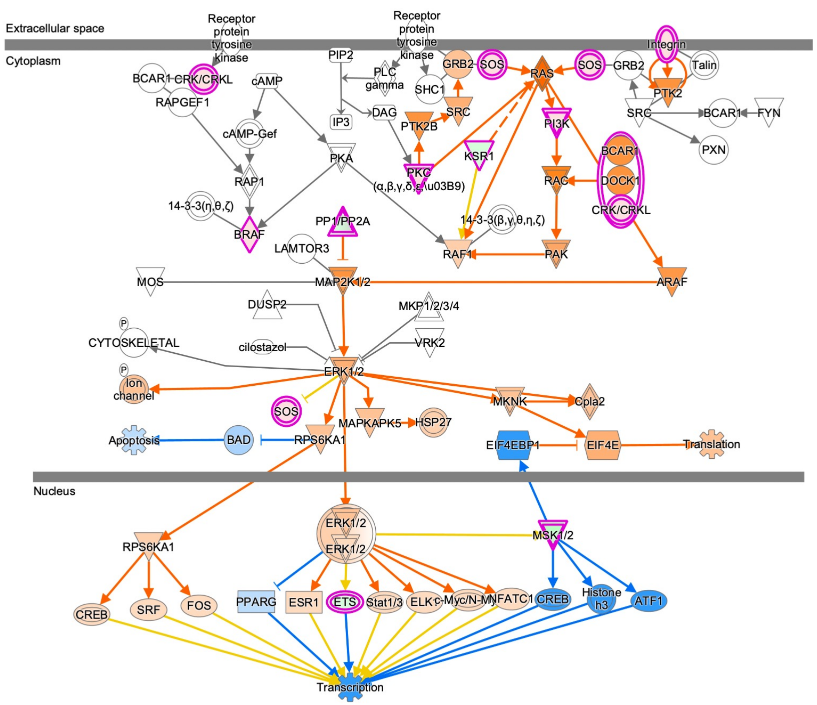


B

#### mTOR signaling

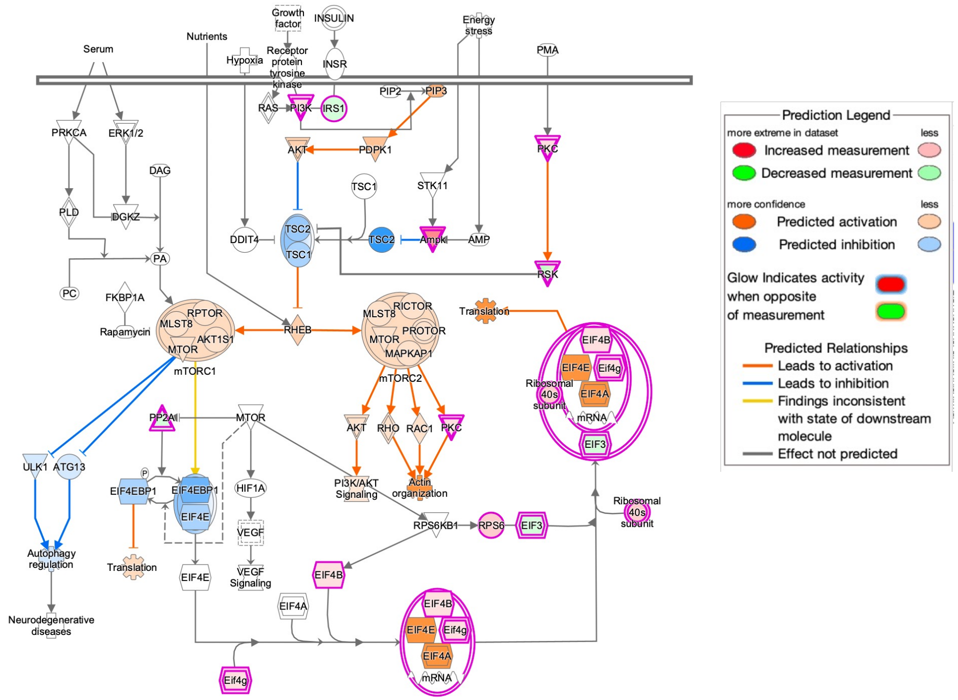

### Supplementary Figures – S5

A

#### Actin cytoskeleton signaling

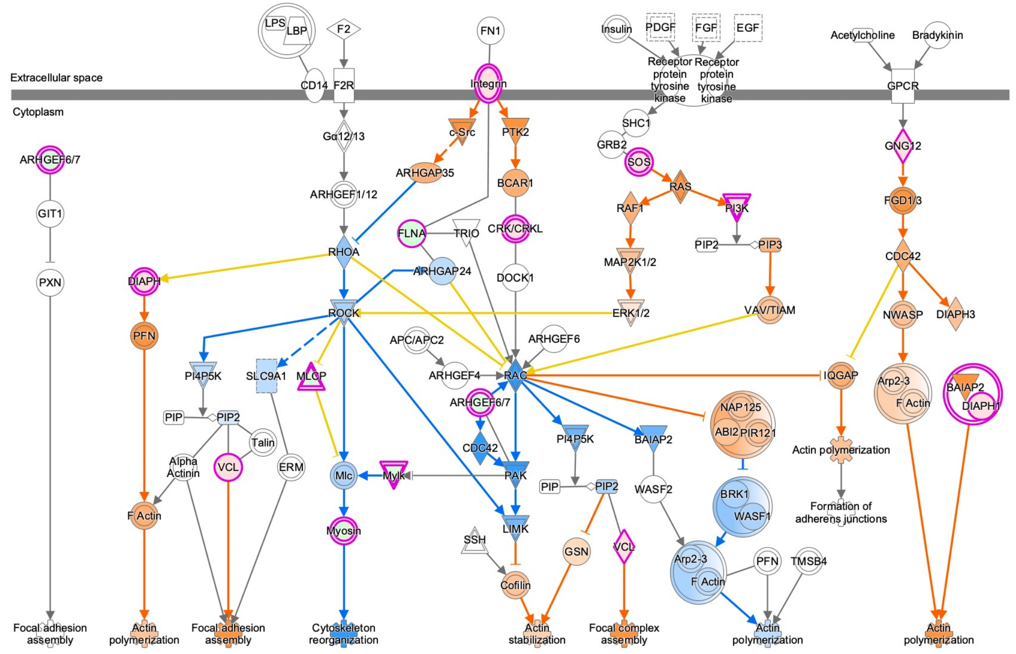


B

#### ILK signaling

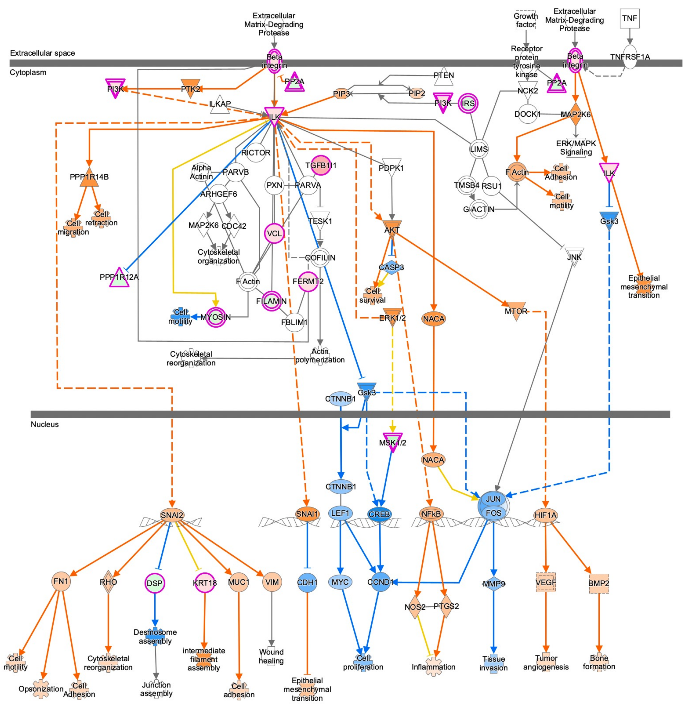


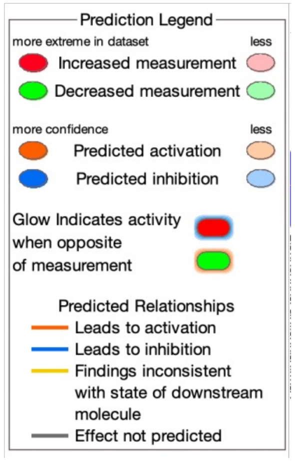


C

#### Integrin signaling

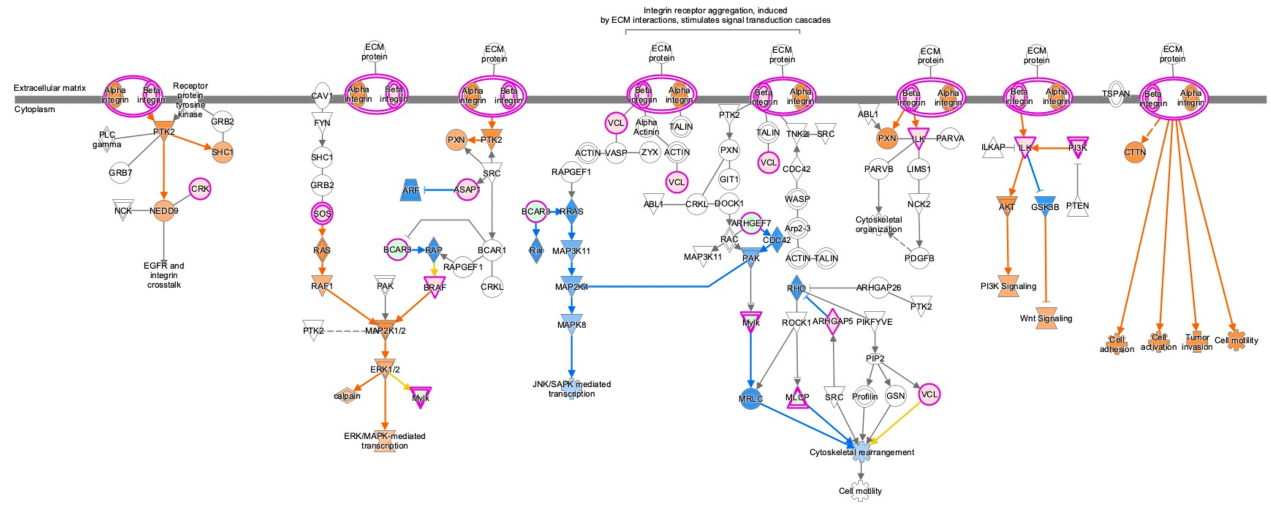

### Supplementary Figures – S6

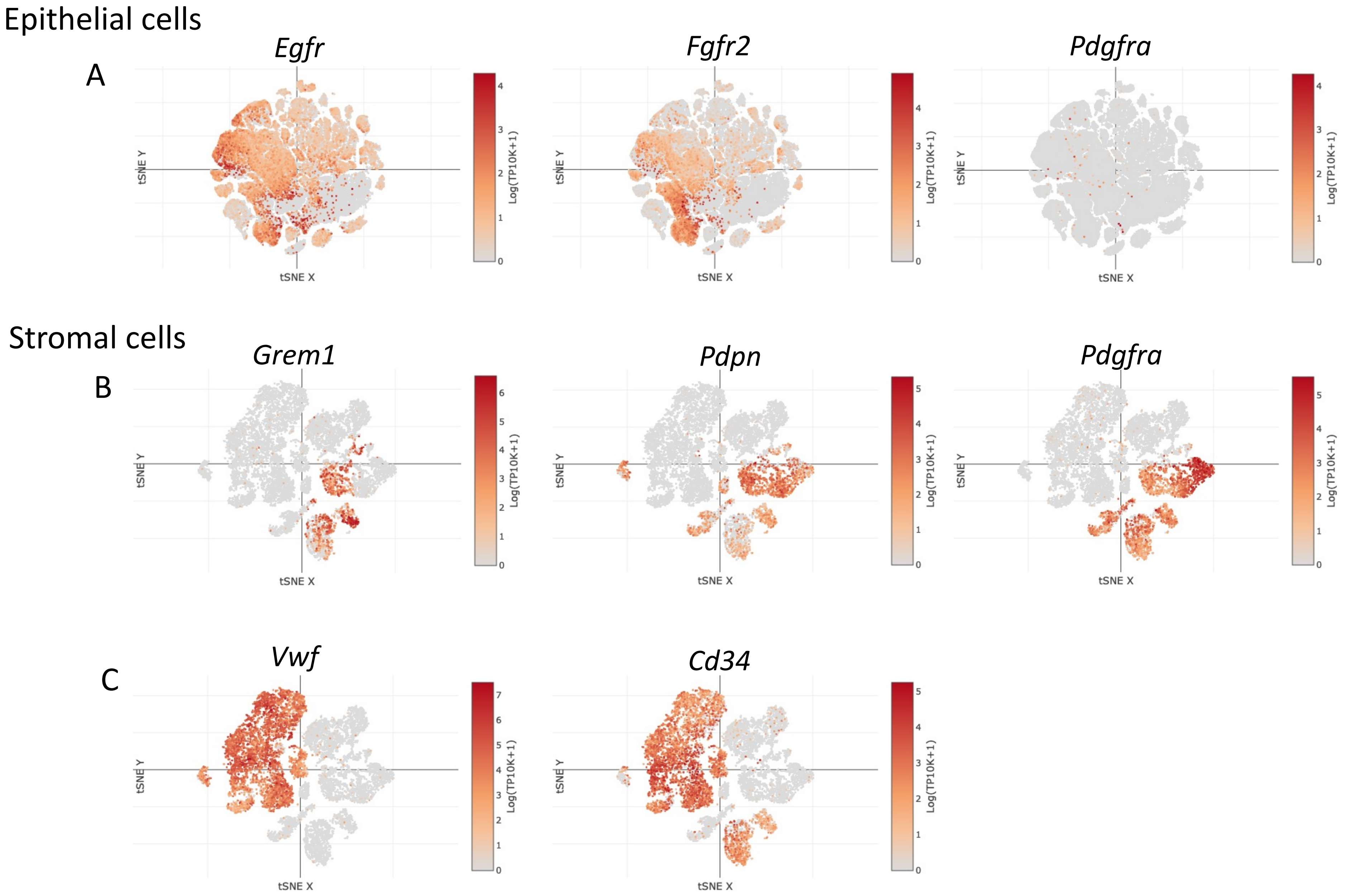

### Supplementary Figures – S7

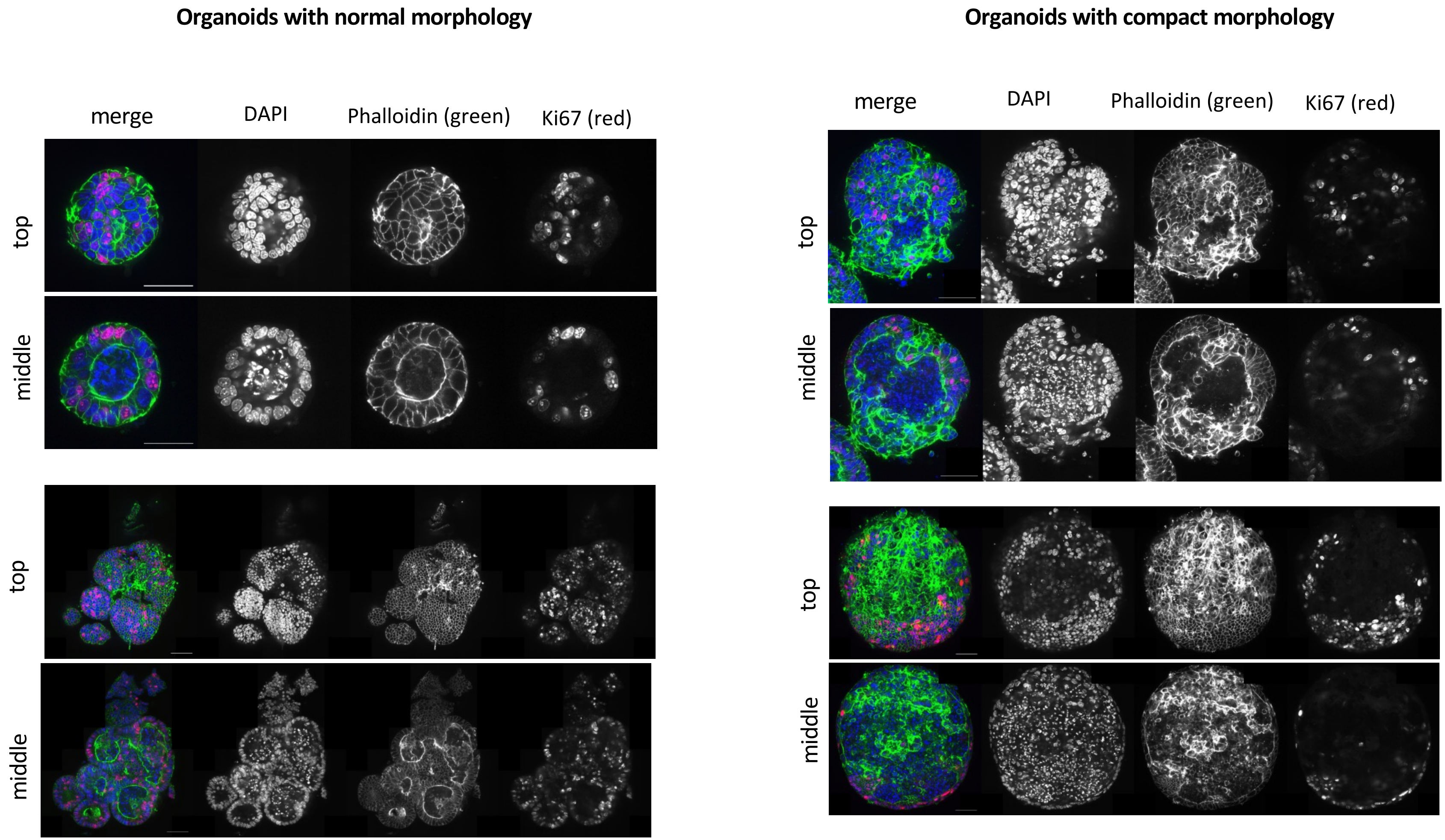

### Supplementary Figures – S8

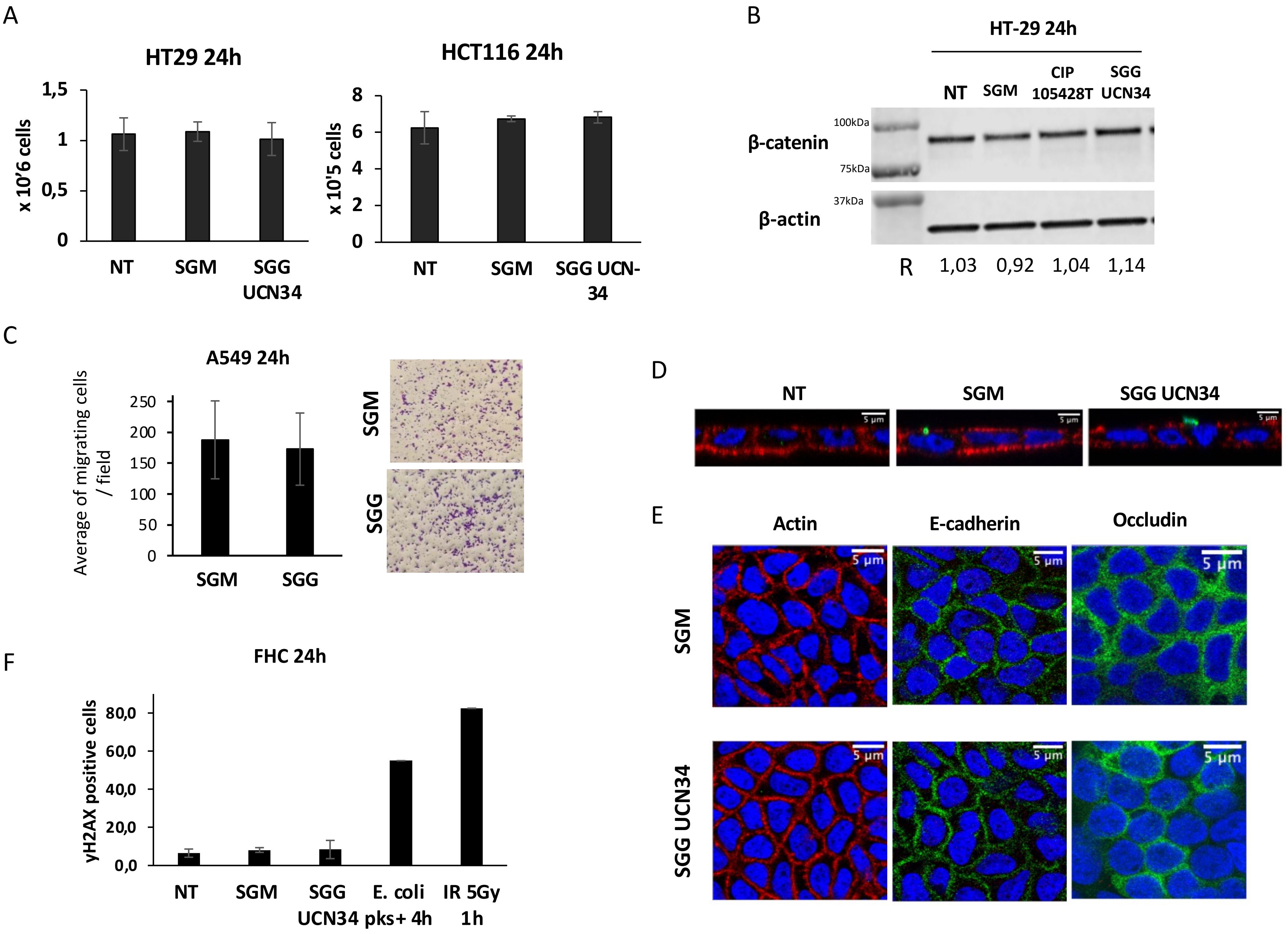

### Supplementary Figures – S9

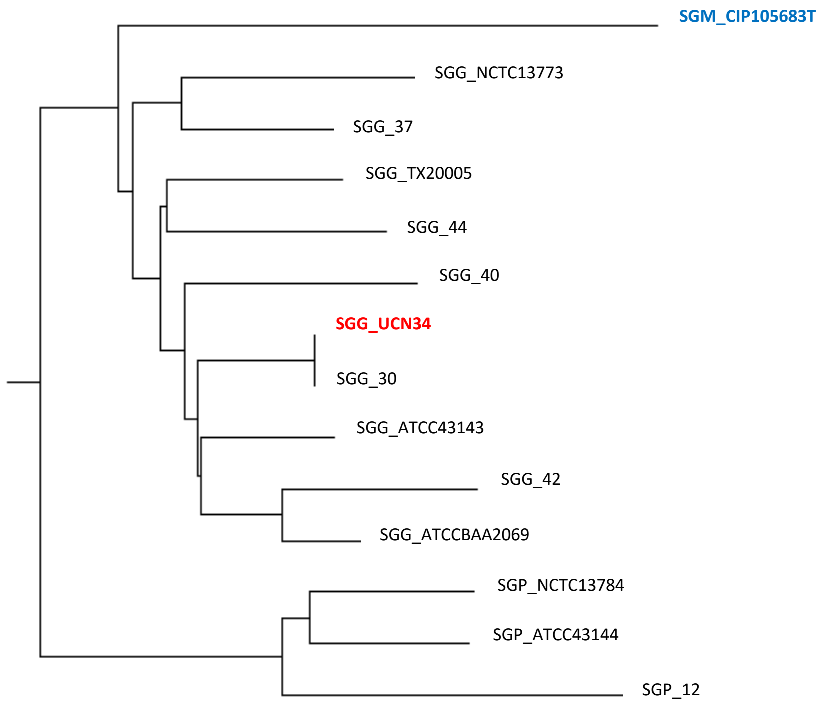
